# The immune system fails to mount a protective response to Gram-positive or Gram-negative bacterial prostatitis

**DOI:** 10.1101/2020.02.29.971051

**Authors:** Federico Lupo, Matthieu Rousseau, Tracy Canton, Molly A. Ingersoll

## Abstract

Bacterial prostatitis affects 1% of men, with increased incidence in the elderly. It is defined by the frequency and urgency to urinate, localized pain, and positive bacterial cultures in expressed seminal fluids. Acute bacterial prostatitis frequently progresses to chronicity, which is marked by recurrent acute episodes interspersed with asymptomatic periods of variable duration. Up to 80% of bacterial prostatitis cases are caused by Gram-negative uropathogenic *E. coli* (UPEC) or Gram-positive *E. faecalis*. Antibiotic treatment is standard of care, however, global dissemination of antimicrobial resistant uropathogens threatens efficacy of therapy. Thus, development of non-antibiotic-based approaches to treat bacterial prostatitis is a priority. One challenge is that the immune response to infection in the prostate is incompletely understood. We used a mouse model of transurethral bacterial instillation to study the immune response to UPEC or *E. faecalis* prostate infection. Both uropathogens exhibited tropism for the prostate over the bladder early post-infection. UPEC infection induced greater proinflammatory cytokine expression and neutrophil and monocyte infiltration compared to *E. faecalis* infection. Following challenge infection, cytokine responses and myeloid cell infiltration were largely comparable to primary infection. Characteristic of memory responses, more lymphoid cells infiltrating the prostate in the second infection compared to the primary infection. Unexpectedly, however, bacterial burden in prostates challenged with either UPEC or *E. faecalis* was equal or greater than in primary infection, despite that an adaptive response to UPEC infection was evident in the bladder of the same animals. Thus, an immune response to primary infection is initiated, however it does not protect against reinfection. Our findings support the idea that chronic or recurrent prostatitis develops in the absence of efficacious immunity to infection. A greater understanding of the mechanisms underlying this observation may point to actionable targets for immunotherapy.

## Introduction

Prostatitis is a common term that encompasses the clinical conditions of acute or chronic bacterial prostatitis, chronic pelvic pain syndrome, and asymptomatic inflammatory prostatitis (Krieger et al., 1999). Characteristic symptoms include urinary frequency and urgency, and suprapubic, lower back, or perianal pain during micturition and sex (Khan et al., 2017). All-cause prostatitis develops in nearly 10% of all men, with increased risks of chronicity after the age of fifty (Krieger et al., 2008; Schaeffer, 2003). Prostatitis is more common in the elderly, similar to urinary tract infection in men (Foxman, 2010; Ingersoll, 2017; Khan et al., 2017; Lipsky, 1999; Rowe and Juthani-Mehta, 2013). An estimated 10% of all prostatitis cases are due to bacterial infection (Khan et al., 2017). Acute bacterial prostatitis is diagnosed by positive bacterial cultures from expressed fluids following prostate massage (Meares and Stamey, 1972; Nickel et al., 2006). Nearly half of patients with acute infection will develop chronic bacterial prostatitis, which is characterized by asymptomatic periods of variable duration interspersed with acute recurrent symptomatic infections (Bowen et al., 2015; Gill and Shoskes, 2016; Khan et al., 2017).

Epidemiological studies from hospitalized clinical cohorts demonstrate that *Escherichia coli* or Gram-positive *Enterococci spp.*, and in particular *E. faecalis,* represent up to 80% of all pathogens isolated from acute or chronic prostatitis patients (Domingue and Hellstrom, 1998; Gill and Shoskes, 2016; Lipsky, 1999; Lipsky et al., 2010). Similar to urinary tract infections (UTI), acute and chronic bacterial prostatitis are typically treated with analgesics to minimize pain, and antibiotics, such as fluoroquinolones or trimethoprim-sulfamethoxazole, to eradicate microorganisms (Gill and Shoskes, 2016; Lipsky, 1999; Lipsky et al., 2010). Depending on the severity of acute symptoms, recommended treatment guidelines may include a single dose of oral fluoroquinolones or intramuscular cephalosporin followed by a ten day tetracycline regimen, or two to four weeks of fluoroquinolones in combination with aminoglycosides (Coker and Dierfeldt, 2016; Lipsky et al., 2010). Chronic prostatitis antibiotic regimens are similar but longer, typically ranging from 6 to 12 weeks (Lipsky et al., 2010). These extended courses are needed to eradicate persistent or recurrent infections due to the reduced penetrance of certain antibiotics (Gill and Shoskes, 2016).

Alarmingly, unusually long regimens of up to 6 months may lead to the loss of efficacy of fluoroquinolones in 10-30% of cases (Naber et al., 2008). These complicated antibiotic regimens highlight the difficulty in treating prostatitis and the challenge to find alternative treatment modalities. The expansion of multidrug-resistant uropathogens further compounds this problem (Stamatiou and Pierris, 2017). To limit development of antibiotic resistance, empirical or “best guess” administration of antibiotics prior to microbiological analysis is recommended only in acute prostatitis (Lipsky et al., 2010). Despite this, most non-bacterial prostatitis cases are treated with antibiotics as this approach, surprisingly, results in amelioration of symptoms (Lipsky et al., 2010). It is unclear whether this effect is ascribable to the antibiotics themselves, to a placebo effect, or whether bacteria are present but undetectable by current testing protocols (Delarosette et al., 1993). Although antibiotic treatment for prostatitis can be challenging, no alternative therapies, including immunomodulatory strategies, exist. One reason for this may be that the immune response provoked during infection is incompletely understood.

Indeed, our current understanding of bacterial colonization, prostate-resident and infiltrating immune cells and their roles in bacterial clearance, or induction of specific memory in the prostate is based on a limited number of studies. For example, with respect to colonization, while the prostatitis-derived *E. coli* CP1 strain expresses fewer virulence factors than the uropathogenic *E. coli* (UPEC) cystitis isolate NU14, it colonizes mouse prostates to a greater extent than NU14 (Rudick et al., 2011), supporting that the ability to colonize the prostate is determined by the pathogen. CP1 colonizes the bladder and the prostate of C57BL/6 mice equally well at 24 hours post-infection (PI), however CP1 persists longer in the prostate compared to the bladder (Rudick et al., 2011). Infection of C57BL/6 mouse prostates with UPEC strain 1667 induces pro-inflammatory cytokine RNA expression, inflammatory cell infiltration, and tissue damage in the prostate over two weeks (Boehm et al., 2012). C3H/HeOuJ mice infected with UPEC 1667 present with persistent inflammation, characterized by collagen deposition (Wong et al., 2014). When treated with a 2 week antibiotic regimen followed by an 8 week washout period, these mice exhibited a reversal in collagen content and diminishment of tissue inflammation despite being infected for 28 days prior to treatment (Wong et al., 2015). When infected for 2-12 months, C57BL/6 mice present with hyperplastic epithelia and greater numbers of macrophages and T_h_17 T cells in the prostate (Simons et al., 2015). Prostate infection with *P. acnes*, a strain associated with benign prostatic hyperplasia, induces inflammation and innate immune cell infiltration only after 1 week of infection (Shinohara et al., 2013). Together, these studies present a somewhat fragmented picture of the innate immune response to bacterial prostatitis and provide little insight into development of adaptive immunity to infectious prostatitis.

We recently postulated that bacterial prostatitis should be classified as a specific type of UTI, similar to cystitis or pyelonephritis (Lupo and Ingersoll, 2019). In addition to sharing associated risks and the route of infection via urethral ascension, UTI and bacterial prostatitis are caused by similar or the same infectious agents, including UPEC and *E. faecalis*. Thus, we hypothesized that the immune response to prostate infection is similar to that observed in the bladder of male mice with cystitis, in which a blunted innate immune response leads to limited immune cell infiltration, but ultimately the development of nonsterilizing specific immunity (Ingersoll et al., 2008; Mora-Bau et al., 2015; Zychlinsky Scharff et al., 2019). To test this hypothesis, we measured colonization and innate and adaptive immunity in a model of acute and chronic prostatitis using Gram-negative UPEC and Gram-positive *E. faecalis* strains. Surprisingly, we observed the response to UPEC infection in the prostate, including robust neutrophil infiltration, was more similar to UTI in female animals than male mice (Ingersoll et al., 2008; Mora-Bau et al., 2015; Zychlinsky Scharff et al., 2019). In both UPEC and *E. faecalis* infection, we found that the prostate exhibited a cytokine signature divergent from that of the bladder and, critically, unlike the bladder, failed to develop protective immunity to challenge infection, providing an explanation for the frequency in which chronic or recurrent prostatitis develops in men.

## Results

### UPEC and E. faecalis display tropism for the mouse prostate

In the course of a previously published study, we observed that, in addition to the bladder, UPEC robustly colonizes the prostate of male mice for up to two weeks (Zychlinsky Scharff et al., 2019). As *E. coli* and *E. faecalis* are the principal uropathogens causing bacterial prostatitis, we wanted to determine the capacity of these bacterial strains to specifically colonize the prostate, in comparison to the bladder, over time. We transurethrally infected 6-8-week old male C57BL/6 mice with 10^7^ CFU of one of two representative uropathogens: UPEC strain UTI89-RFP-kan^R^, resistant to kanamycin and expressing red fluorescent protein, or *E. faecalis* strain OG1RF, resistant to rifampicin (Dunny et al., 1978; Mora-Bau et al., 2015). We quantified bacterial burden in homogenized bladders and prostates at 1, 2, and 14 days PI, to model acute and chronic infection. We observed that while UPEC robustly colonized both organs in male mice, UPEC bacterial burden was significantly higher in prostates, compared to bladders, of infected mice at day 1 (**Figure 1A**). Differences in UPEC colonization between prostates and bladders were no longer apparent at 2 or 14 days PI. We reported that male mouse bladders remain colonized up to 28 days following UPEC infection (Zychlinsky Scharff et al., 2019). Here, we observed that prostates and bladders were still colonized with UPEC at 42 days PI (**Figure S1A**). Whereas UPEC colonized the prostate and bladder robustly, *E. faecalis* bacterial burden was lower in the bladder, and in line with previous reports of *E. faecalis* burden in infected female mice (**Figure 1B**) (Kau et al., 2005; Singh et al., 2007; Tien et al., 2017). Notably, *E. faecalis* displayed a more consistent infection and pronounced tropism for the prostate over the bladder at day 1, 2, and 14 PI (**Figure 1B**). Finally, as polymicrobial infections are common in humans and may lead to complicated UTI or urosepsis (Brogden et al., 2004; Kline and Lewis, 2016; Siegmanigra et al., 1993; Siegmanigra et al., 1994; Tay et al., 2016), we measured bacterial burdens in bladders and prostates following co-infection with UPEC and *E. faecalis*. We instilled a total of 10^7^ CFU in an approximate 1:1 ratio of each strain, and observed that UPEC bacterial burden was significantly higher in prostates compared to the bladder at day 1 and 2 PI (**Figure 1C**). In the context of this co-infection, *E. faecalis* colonized both organs, however, statistically significant differences in bacterial burden were no longer apparent between the bladder and the prostate due to the increased variance in *E. faecalis* CFU in the bladder at day 1 and 2 PI (**Figure 1D**). Having established mono and polymicrobial prostate infection models with two common uropathogens, we used these models to investigate the immune response to prostatitis, with a focus on the early events following both primary and challenge infection, given the frequency in which these infections recur or become chronic.

**Figure 1.**
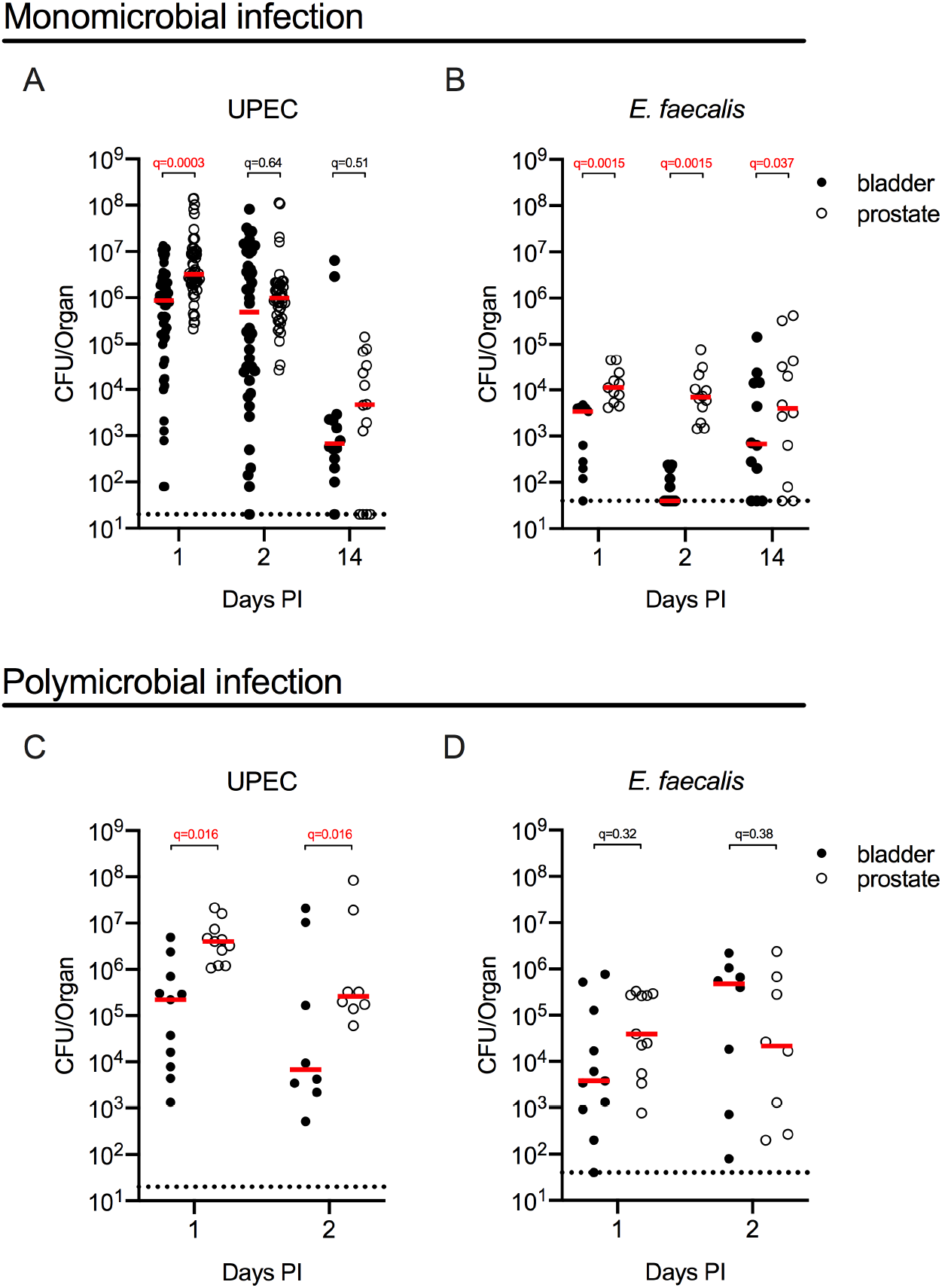
UPEC and *E. faecalis* display tropism for the prostate over the bladder. 6-8-week old male C57BL/6 mice were infected with 10^7^ CFU of (**A**) UPEC strain UTI89-RFP-kan^R^, (**B**) *E. faecalis* strain OG1RF, or (**C-D**) a total of 10^7^ CFU of both bacteria strains in an approximate 1:1 ratio. Graphs depict the CFU/organ at the indicated day post-infection (PI). Each circle represents one mouse, filled circles represent bladder CFU, open circles represent prostate CFU. Red lines indicate the median value of each group and black dotted lines represent the limit of detection of the assay, 20 or 40 CFU/organ. Graphs show paired data as one bladder and one prostate were collected from each mouse. Data are pooled from 2-10 experiments, with 4-7 mice per group in each experiment. CFU in the bladder and prostate were compared at each time point using the non-parametric Wilcoxon test for paired data and *p*-values were corrected for multiple testing within each infection scenario (UPEC, *E. faecalis*, polymicrobial) using the false discovery rate (FDR) method. *q*-values meeting the criteria for statistical significance (*q* <0.05) are depicted in red.

### The innate immune prostate cytokine signature is divergent from the bladder during infection

Having established that both UPEC and *E. faecalis* robustly infects the prostate, we next considered the inflammatory response elicited by bacterial colonization. As we hypothesized that the response to uropathogen infection in the prostate is similar to that observed in the bladder in male C57BL/6 mice, we included analysis of this organ for comparison. We infected male C57BL/6 mice with 10^7^ CFU of UTI89-RFP-kan^R^, OG1RF, or a 1:1 ratio of both strains and analyzed the expression of 13 cytokines by bead array in prostate and bladder homogenates at day 1 and 2 days PI. UPEC-infected prostates expressed statistically significantly higher levels of IL-1α, IL-6, CCL2, IL-1β, IL-17A, and TNF-α compared to bladder tissue at 1 and 2 days PI (**Figure 2A, Table 1**). IL-23 levels were significantly elevated in the prostate over the bladder only at 2 days PI (**Table 1**). Interestingly, IL-6 and IL-1β levels in naïve prostate were significantly higher than in naive bladders, although the reasons for this are unclear (**Figure 2A, Table 1**). When considering the innate response within the prostate, we observed that in UPEC infection IL-1α, IL-6, CCL2, IL-1β, IL-17A, and TNF-α were all elevated over levels in naïve tissue 1 day PI (**Figure 2A**, **Table 2**). IL-1α, CCL2, and TNF-α remained significantly elevated compared to naïve tissue 2 days PI, however, the overall response was short-lived as we measured a significant reduction in IL-27, IL-23, IL-6, CCL2, IL-1β, IL-17A, TNF-α, and CSF2 from day 1 to day 2 PI (**Figure 2A, Table 2**). Finally, despite persistent infection, evidenced by consistent bacteriuria, cytokine levels in UPEC-infected tissues 42 days PI were not different than those measured in naïve tissue, with the exception of TNFα (**Figure S1B-C**).

**Figure 2.**
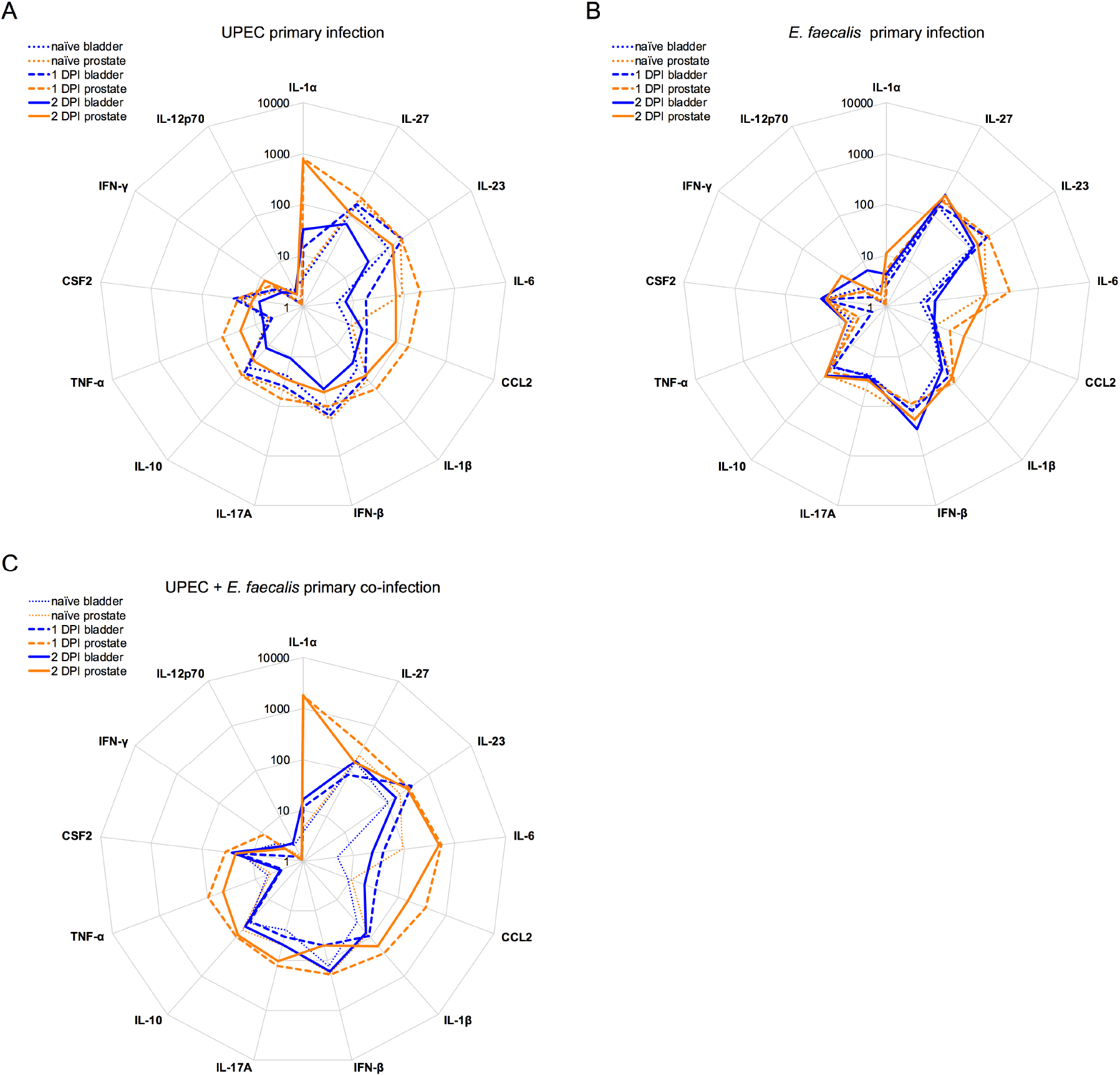
The cytokine response to prostate infection diverges from that observed in the bladder. 6-8-week old male C57BL/6 mice were infected with 10^7^ CFU of (**A**) UPEC strain UTI89-RFP-kan^R^, (**B**) *E. faecalis* strain OG1RF, or (**C**) a total of 10^7^ CFU of both bacteria strains in an approximate 1:1 ratio and sacrificed 1 or 2 days PI. Naïve mice were included for baseline determination. Thirteen cytokines were measured in the supernatants of homogenized bladders and prostates by multiplex analysis. Spider plots show the median values of absolute cytokine levels (pg/mL) on a log-scale in naïve (dotted lines) or infected animals at 24 hours (dashed lines) and 48 hours post-infection (PI) (solid lines) in which prostate values are depicted in orange and bladder values are shown in blue. Data are pooled from 2-7 experiments, with 3-6 mice per group in each experiment. DPI - days post-infection. Significance was determined using the Mann-Whitney nonparametric test with correction for multiple testing to determine the false discovery rate (FDR) adjusted p value: q<0.05. All *p*- and *q*-values for each comparison are listed in **Tables 1-6**.

**Table 1:** Statistical testing of analytes defining the innate immune response to UPEC infection in the bladder versus the prostate.

| *Analyte | Bladder vs prostate naïve |  | Bladder vs prostate, 1 DPI |  | Bladder vs prostate, 2 DPI |  |
| --- | --- | --- | --- | --- | --- | --- |
|  | *p*-value | *q*-value | *p*-value | *q*-value | *p*-value | *q*-value |
| IL-1α | 2.33E-01 | 6.54E-01 | **1.81E-12** | **1.18E-11** | **6.84E-03** | **1.48E-02** |
| IL-27 | 7.40E-01 | 9.77E-01 | 3.63E-01 | 4.29E-01 | 4.17E-01 | 4.93E-01 |
| IL-23 | 5.84E-01 | 9.49E-01 | 5.36E-01 | 5.36E-01 | **4.43E-03** | **1.44E-02** |
| IL-6 | **1.79E-05** | **2.32E-04** | **1.32E-09** | **4.28E-09** | **4.31E-04** | **5.60E-03** |
| CCL2 | 5.69E-01 | 9.49E-01 | **2.39E-10** | **1.03E-09** | **1.50E-03** | **7.28E-03** |
| IL-1β | **7.10E-03** | **4.62E-02** | **3.87E-05** | **1.01E-04** | **6.84E-03** | **1.48E-02** |
| IFN-β | 9.12E-01 | 9.88E-01 | 2.40E-01 | 3.47E-01 | 7.81E-01 | 7.81E-01 |
| IL-17A | 7.52E-01 | 9.77E-01 | **1.15E-02** | **2.49E-02** | **1.13E-02** | **2.10E-02** |
| IL-10 | 2.52E-01 | 6.54E-01 | 5.19E-01 | 5.36E-01 | 1.11E-01 | 1.80E-01 |
| TNF-α | 8.59E-01 | 9.88E-01 | **3.79E-13** | **4.93E-12** | **1.68E-03** | **7.28E-03** |
| CSF2 | 1.00E+00 | 1.00E+00 | 3.03E-01 | 3.94E-01 | 4.17E-01 | 4.93E-01 |
| IFN-γ | 5.05E-01 | 9.49E-01 | **4.27E-02** | 7.93E-02 | 3.19E-01 | 4.61E-01 |
| IL-12p70 | 2.20E-01 | 6.54E-01 | 1.52E-01 | 2.46E-01 | 4.67E-01 | 5.06E-01 |

\*Values are ordered according to expression level of measured cytokines in UPEC-infected prostates at 1 day PI (DPI). *p*-values were determined by Mann-Whitney nonparametric test, and corrected for multiple testing to determine *q*-values (false discovery rate adjusted *p*-value). *p*-values and *q*-values <0.05 are shown in bold.

**Table 2:**
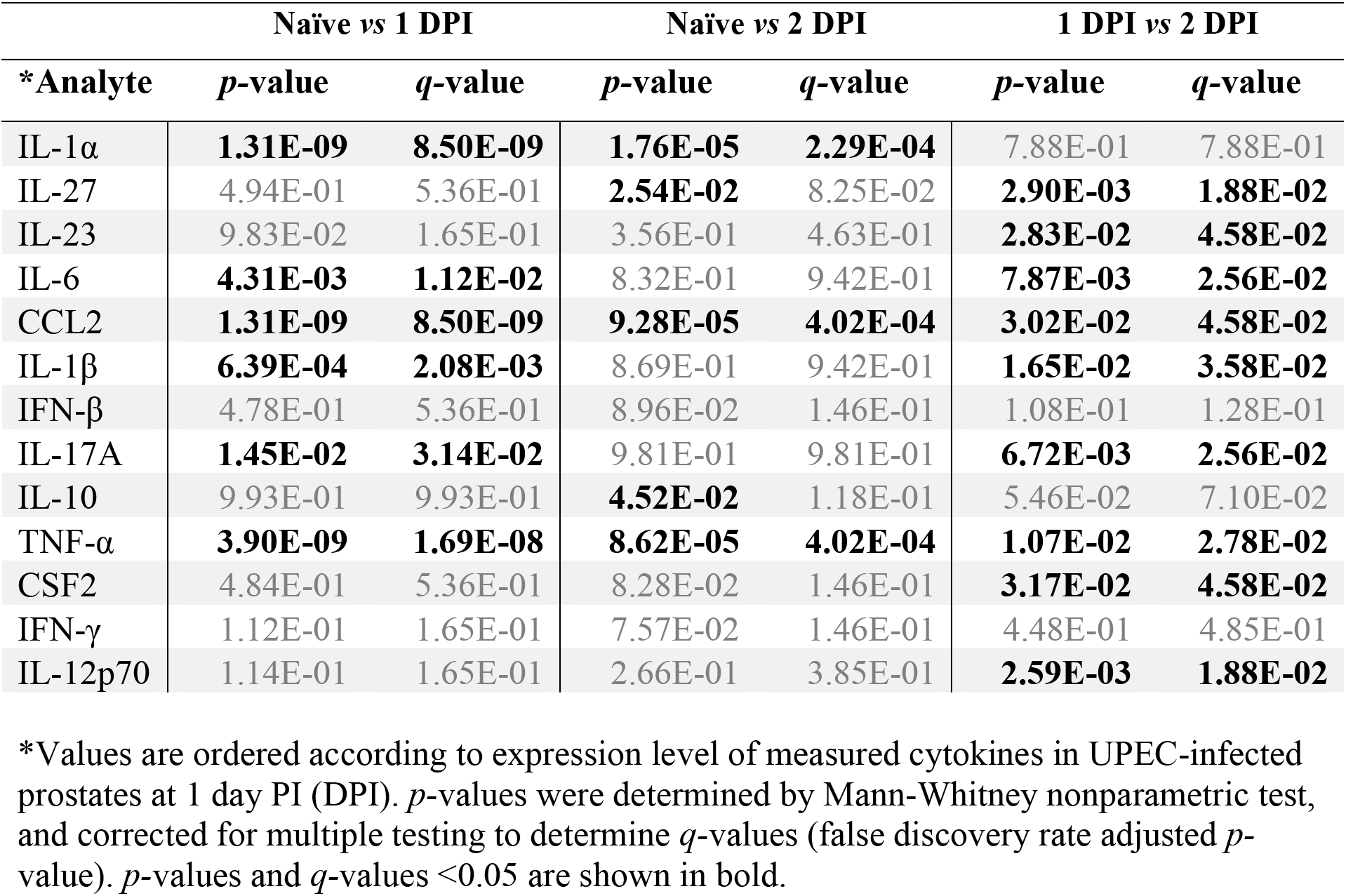
Statistical testing of analytes defining the innate immune response to UPEC infection in the prostate over time.

In *E. faecalis* infected mice, IL-1α, IL-6, CCL2, IL-1β, and TNF-α were expressed at levels significantly higher in the prostate than in the bladder at 1 day PI, and IL-1α, IL-6, CCL2, and IL-1β remained elevated at 2 days PI (**Figure 2B, Table 3**). Notably, no cytokines were significantly elevated over naïve tissue levels in the prostate at 1 day PI with *E. faecalis* after correction for multiple testing (**Figure 2B**, **Table 4**). At 2 days PI with *E. faecalis*, IL-1α, CCL2, and IFN-**γ** were elevated over naïve tissue levels, and similar to that seen in UPEC infection, the response contracted from 1 to 2 days PI, with a reduction in IL-23 and IL-6. By contrast, TNF-α and IFN-**γ** were expressed at significantly higher levels at 2 days PI compared to day 1 PI (**Figure 2B, Table 4**). Globally, cytokine levels induced after *E. faecalis* infection were lower than those measured in UPEC-infected bladders and prostates, closely mirroring that measured in naïve tissue (**Figure 2A, B**).

**Table 3:**
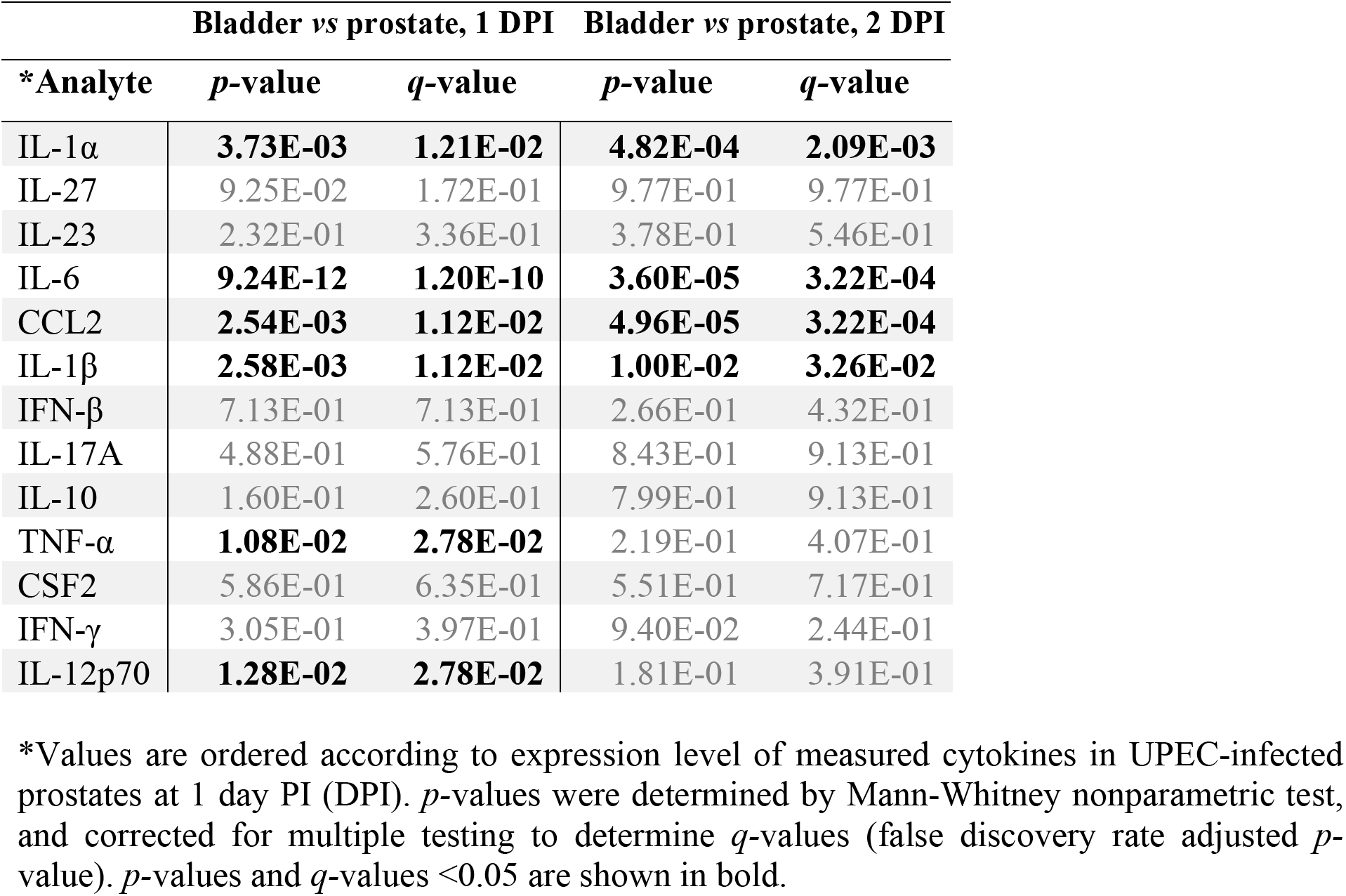
Statistical testing of analytes defining the innate immune response to *E. faecalis* infection in the bladder versus the prostate.

| *Analyte | Bladder vs prostate, 1 DPI |  | Bladder vs prostate, 2 DPI |  |
| --- | --- | --- | --- | --- |
|  | *p*-value | *q*-value | *p*-value | *q*-value |
| IL-1α | **3.73E-03** | **1.21E-02** | **4.82E-04** | **2.09E-03** |
| IL-27 | 9.25E-02 | 1.72E-01 | 9.77E-01 | 9.77E-01 |
| IL-23 | 2.32E-01 | 3.36E-01 | 3.78E-01 | 5.46E-01 |
| IL-6 | **9.24E-12** | **1.20E-10** | **3.60E-05** | **3.22E-04** |
| CCL2 | **2.54E-03** | **1.12E-02** | **4.96E-05** | **3.22E-04** |
| IL-1β | **2.58E-03** | **1.12E-02** | **1.00E-02** | **3.26E-02** |
| IFN-β | 7.13E-01 | 7.13E-01 | 2.66E-01 | 4.32E-01 |
| IL-17A | 4.88E-01 | 5.76E-01 | 8.43E-01 | 9.13E-01 |
| IL-10 | 1.60E-01 | 2.60E-01 | 7.99E-01 | 9.13E-01 |
| TNF-α | **1.08E-02** | **2.78E-02** | 2.19E-01 | 4.07E-01 |
| CSF2 | 5.86E-01 | 6.35E-01 | 5.51E-01 | 7.17E-01 |
| IFN-γ | 3.05E-01 | 3.97E-01 | 9.40E-02 | 2.44E-01 |
| IL-12p70 | **1.28E-02** | **2.78E-02** | 1.81E-01 | 3.91E-01 |

\*Values are ordered according to expression level of measured cytokines in UPEC-infected prostates at 1 day PI (DPI). *p*-values were determined by Mann-Whitney nonparametric test, and corrected for multiple testing to determine *q*-values (false discovery rate adjusted *p*-value). *p*-values and *q*-values <0.05 are shown in bold.

**Table 4:** Statistical testing of analytes defining the innate immune response to E. faecalis infection in the prostate over time.

| *Analyte | Naïve vs 1 DPI |  | Naïve vs 2 DPI |  | 1 DPI vs 2 DPI |  |
| --- | --- | --- | --- | --- | --- | --- |
|  | *p*-value | *q*-value | *p*-value | *q*-value | *p*-value | *q*-value |
| IL-1α | 3.29E-01 | 6.31E-01 | **2.28E-03** | **1.48E-02** | **4.90E-02** | 9.11E-02 |
| IL-27 | 9.70E-01 | 9.70E-01 | 1.00E+00 | 1.00E+00 | 7.32E-01 | 7.62E-01 |
| IL-23 | **1.09E-02** | 6.88E-02 | 4.65E-01 | 8.63E-01 | **1.06E-02** | **3.43E-02** |
| IL-6 | **1.59E-02** | 6.88E-02 | 7.87E-01 | 9.68E-01 | **1.74E-02** | **4.52E-02** |
| CCL2 | **1.31E-02** | 6.88E-02 | **6.53E-04** | **8.49E-03** | 8.32E-02 | 1.35E-01 |
| IL-1β | 4.37E-01 | 6.31E-01 | 8.19E-01 | 9.68E-01 | 3.08E-01 | 4.45E-01 |
| IFN-β | 2.90E-01 | 6.31E-01 | 6.57E-01 | 9.48E-01 | **4.90E-02** | 9.11E-02 |
| IL-17A | 7.21E-01 | 9.13E-01 | 3.85E-01 | 8.35E-01 | 4.68E-01 | 6.08E-01 |
| IL-10 | 3.98E-01 | 6.31E-01 | 6.32E-01 | 9.48E-01 | 5.82E-01 | 6.87E-01 |
| TNF-α | 3.67E-01 | 6.31E-01 | **1.87E-02** | 6.09E-02 | **8.00E-03** | **3.43E-02** |
| CSF2 | 9.27E-01 | 9.70E-01 | 9.49E-01 | 1.00E+00 | 7.62E-01 | 7.62E-01 |
| IFN-γ | 7.72E-01 | 9.13E-01 | **3.95E-03** | **1.71E-02** | **9.71E-04** | **6.31E-03** |
| IL-12p70 | **3.58E-02** | 1.16E-01 | 2.94E-01 | 7.63E-01 | **1.02E-04** | **1.33E-03** |

\*Values are ordered according to expression level of measured cytokines in UPEC-infected prostates at 1 day PI (DPI). *p*-values were determined by Mann-Whitney nonparametric test, and corrected for multiple testing to determine *q*-values (false discovery rate adjusted *p*-value). *p*-values and *q*-values <0.05 are shown in bold.

In the co-infection scenario, more cytokines were significantly elevated in the prostate compared to the bladder than in either monomicrobial infection at day 1 PI, including IL-1α, IL-27, IL-6, CCL2, IL-1β, IL-17A, IL-10, TNF-α, CSF2, and IFN-**γ** (**Figure 2C, Table 5**). IL-1α, IL-6, IL-1β, and TNF-α levels in the prostate remained elevated over bladder-associated levels at 2 days PI (**Figure 2C, Table 5**). With the exception of IL-27 and IFN-β, all cytokines measured were significantly increased at 1 day PI compared to naïve levels (**Figure 2C, Table 6**). While IL-1α, IL-6, CCL2, IL-1β, IL-17A, and TNF-α remained elevated compared to naïve tissue at 2 days PI, no cytokines were significantly different between 1 and 2 days PI in the polymicrobial prostate infection (**Figure 2C, Table 6**). Finally, although IL-12p70 levels were statistically significantly different in several scenarios, such as between day 1 and day 2 PI in UPEC and *E. faecalis* infection, expression levels were near or at the limit of detection in all infection scenarios.

**Table 5:** Statistical testing of analytes defining the innate immune response to polymicrobial infection in the bladder versus the prostate.

| *Analyte | Bladder vs prostate, 1 DPI |  | Bladder vs prostate, 2 DPI |  |
| --- | --- | --- | --- | --- |
|  | *p*-value | *q*-value | *p*-value | *q*-value |
| IL-1α | **4.62E-12** | **3.00E-11** | **1.55E-04** | **2.02E-03** |
| IL-27 | **3.11E-03** | **4.49E-03** | 1.00E+00 | 1.00E+00 |
| IL-23 | 4.20E-01 | 4.20E-01 | 6.50E-02 | 1.41E-01 |
| IL-6 | **1.07E-06** | **3.49E-06** | **1.33E-02** | **4.33E-02** |
| CCL2 | **2.08E-06** | **5.41E-06** | **4.99E-02** | 1.30E-01 |
| IL-1β | **4.86E-13** | **6.32E-12** | **6.99E-03** | **3.03E-02** |
| IFN-β | **4.97E-02** | 5.87E-02 | 7.01E-01 | 9.12E-01 |
| IL-17A | **2.80E-05** | **6.07E-05** | 4.42E-01 | 7.18E-01 |
| IL-10 | **8.73E-04** | **1.42E-03** | 5.05E-01 | 7.30E-01 |
| TNF-α | **6.39E-09** | **2.77E-08** | **2.68E-03** | **1.74E-02** |
| CSF2 | **2.57E-02** | **3.35E-02** | 9.59E-01 | 1.00E+00 |
| IFN-γ | **6.86E-05** | **1.27E-04** | 1.00E+00 | 1.00E+00 |
| IL-12p70 | 8.68E-02 | 9.40E-02 | 8.47E-02 | 1.57E-01 |

\*Values are ordered according to expression level of measured cytokines in UPEC-infected prostates at 1 day PI (DPI). *p*-values were determined by Mann-Whitney nonparametric test, and corrected for multiple testing to determine *q*-values (false discovery rate adjusted *p*-value). *p*-values and *q*-values <0.05 are shown in bold.

**Table 6:** Statistical testing of analytes defining the innate immune response to polymicrobial infection in the prostate over time.

| *Analyte | Naïve vs 1 DPI |  | Naïve vs 2 DPI |  | 1 DPI vs 2 DPI |  |
| --- | --- | --- | --- | --- | --- | --- |
|  | *p*-value | *q*-value | *p*-value | *q*-value | *p*-value | *q*-value |
| IL-1α | **5.77E-08** | **1.50E-07** | **7.04E-05** | **2.23E-04** | 7.07E-01 | 8.02E-01 |
| IL-27 | 1.28E-01 | 1.38E-01 | 6.77E-01 | 7.33E-01 | **3.81E-02** | 9.91E-02 |
| IL-23 | **1.76E-04** | **3.27E-04** | **4.70E-02** | 7.64E-02 | 7.40E-01 | 8.02E-01 |
| IL-6 | **6.63E-10** | **4.31E-09** | **8.96E-06** | **1.16E-04** | 2.75E-01 | 3.97E-01 |
| CCL2 | **5.79E-08** | **1.50E-07** | **7.08E-05** | **2.23E-04** | **2.32E-02** | 7.54E-02 |
| IL-1β | **6.93E-11** | **9.00E-10** | **8.58E-05** | **2.23E-04** | **1.78E-02** | 7.54E-02 |
| IFN-β | 7.33E-01 | 7.33E-01 | 3.03E-01 | 3.94E-01 | 5.20E-02 | 1.13E-01 |
| IL-17A | **2.12E-06** | **4.58E-06** | **3.53E-03** | **7.65E-03** | 1.21E-01 | 1.97E-01 |
| IL-10 | **4.04E-02** | **4.78E-02** | 1.44E-01 | 2.07E-01 | 5.42E-01 | 7.05E-01 |
| TNF-α | **5.54E-08** | **1.50E-07** | **8.14E-05** | **2.23E-04** | **1.17E-02** | 7.54E-02 |
| CSF2 | **4.01E-03** | **5.22E-03** | 4.20E-01 | 4.97E-01 | 9.47E-02 | 1.76E-01 |
| IFN-γ | **4.01E-03** | **5.22E-03** | 8.24E-01 | 8.24E-01 | **2.26E-02** | 7.54E-02 |
| IL-12p70 | **4.12E-04** | **6.69E-04** | **3.42E-02** | 6.35E-02 | 8.60E-01 | 8.60E-01 |

\*Values are ordered according to expression level of measured cytokines in UPEC-infected prostates at 1 day PI (DPI). *p*-values were determined by Mann-Whitney nonparametric test, and corrected for multiple testing to determine *q*-values (false discovery rate adjusted *p*-value). *p*-values and *q*-values <0.05 are shown in bold.

### Cytokine levels following challenge are reduced in polymicrobial primary infection

In the case of UPEC infection, although prostates remained colonized up to 42 days, cytokine levels quickly returned to naïve levels. To determine whether persistent infection, in general, rendered animals refractory to induction of inflammatory mediators in the case of a second or challenge infection, we measured cytokine concentrations in the supernatant from tissue homogenates of challenged prostates. We infected cohorts of male C57BL/6 mice with 10^7^ CFU of UPEC, OG1RF, or a 1:1 ratio of both strains and monitored bacterial clearance by culturing urine samples every 2-3 days over the course of one month. Following antibiotic treatment to resolve infection and a short washout period, mice with sterile urine were challenged with a second infection. Mice that received UPEC or *E. faecalis* were challenged with an isogenic strain of the same species used for primary infection. Animals receiving a primary polymicrobial infection were monomicrobially challenged with either UPEC or *E. faecalis.* Cytokine concentrations of IL-1α, IL-17A, TNF-α, and IFN-γ were significantly elevated 1 day PI in UPEC-challenged prostates that had previously been exposed to a primary UPEC infection, whereas IL-27 and IFN-β were significantly decreased following challenge infection (**Figure 3A**, **Table 7**). By contrast, *E. faecalis* challenge infection induced increased expression only of IL-1α and CCL2 over levels observed following primary infection (**Figure 3B**, **Table 7**). IL-12p70 also was significantly different between primary and challenge infection with UPEC or *E. faecalis,* however, as in primary infections, expression levels of this cytokine were very low. Surprisingly, total cytokine expression was significantly reduced following challenge with either UPEC or *E. faecalis* compared to the response induced by primary polymicrobial infection (**Figure 3C**, **Table 8**). Indeed, IL-1α, IL-6, CCL2, IL-1β, IL-17A, IL-10, TNF-α, and CSF2, were all significantly reduced after UPEC infection, and all cytokines measured except for IL-23 were reduced following *E. faecalis* infection (**Figure 3C**, **Table 8**). Notably, we did not observe this pattern in the bladder of these same animals (**Figure S2)**.

**Figure 3.**
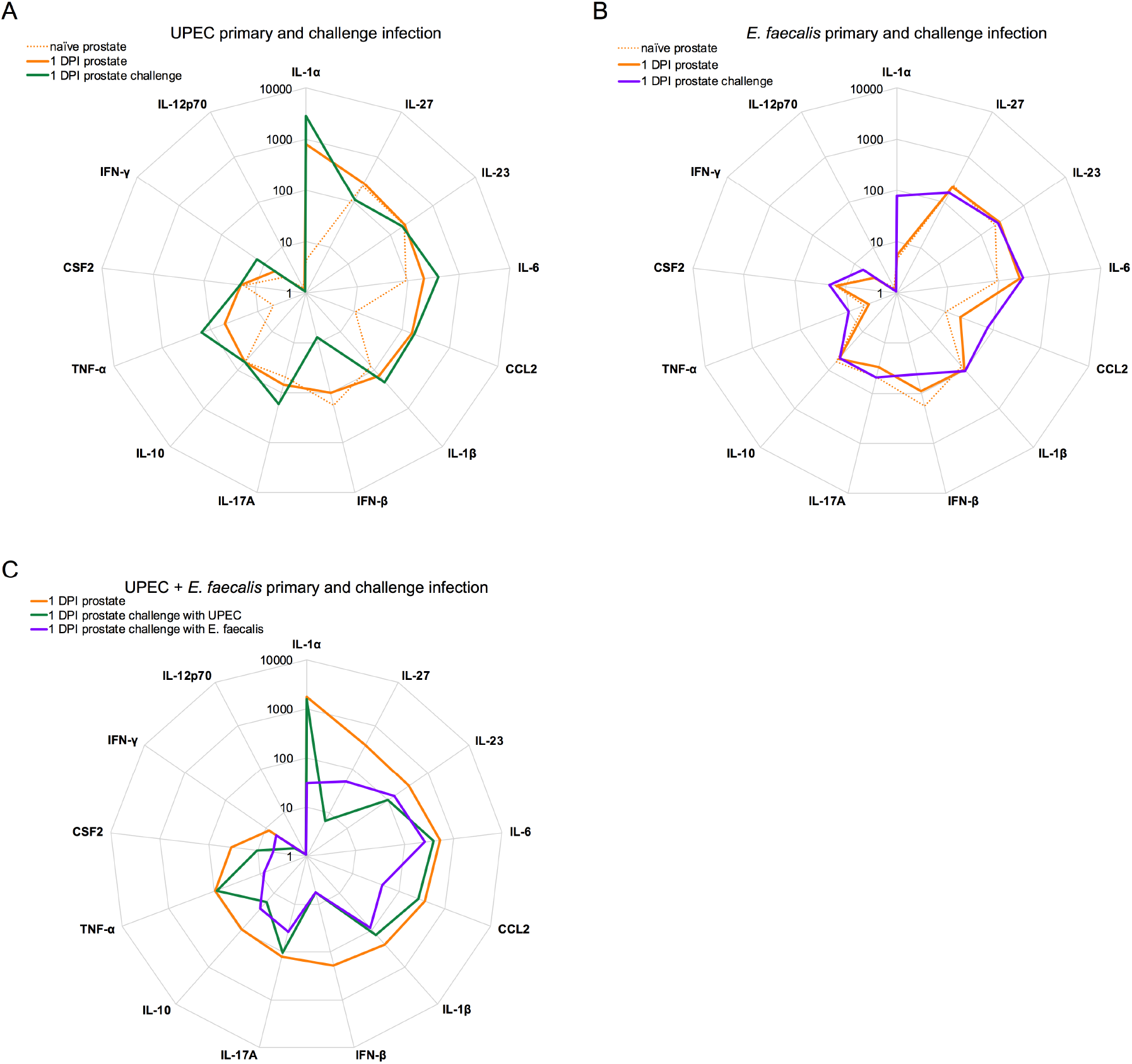
The cytokine response is largely unchanged following challenge infection. 6-8-week old male C57BL/6 mice were infected with 10^7^ CFU of (**A**) UPEC strain UTI89-RFP-kan^R^, (**B**) *E. faecalis* strain OG1RF, or (**C**) a total of 10^7^ CFU of both bacteria strains in an approximate 1:1 ratio. At 25-30 days post-infection (PI), all animals were treated with one or two cycles of antibiotics as described in the Materials and methods for 5 days, followed by a 3-5 day washout period. Mice with sterile urine were challenged with 10^7^ CFU of (**A**) UPEC strain UTI89-GFP-amp^R^, (**B**) *E. faecalis* strain OG1RF_intergenicRS00490RS00495::Tn, or (**C**) in primary polmicrobially-infected mice, half of the cohort was challenged with UTI89-GFP-amp^R^ and the other half was challenged with OG1RF_intergenicRS00490RS00495::Tn. Thirteen cytokines were measured in the supernatants of prostate homogenates by multiplex assay. Spider plots show the median values of absolute prostate cytokine levels (pg/mL) on a log-scale at 24 hours post-primary (dotted lines) or challenge infection (solid lines). DPI - days post-infection. Data are pooled from 2-7 experiments, with 2-6 mice per group in each experiment. In **A, B**, values for prostate cytokine expression in naïve and primary infected prostate are replotted (dotted lines) from Figure 2 for ease in comparing primary to challenge infection. Significance was determined using the Mann-Whitney nonparametric test with correction for multiple testing to determine the false discovery rate (FDR) adjusted p value: q<0.05. All *p*- and *q*-values for each comparison are listed in **Tables 7-8**.

**Table 7:** Statistical testing of analytes expressed during monomicrobial primary or challenge infection with UPEC or *E. faecalis*.

| *Analyte | Primary vs challenge,<br>UPEC |  | Primary vs challenge,<br>*E. faecalis* |  |
| --- | --- | --- | --- | --- |
|  | *p*-value | *q*-value | *p*-value | *q*-value |
| IL-1α | **3.47E-04** | **3.83E-03** | **8.69E-04** | **5.65E-03** |
| IL-27 | **5.54E-03** | **1.44E-02** | 2.44E-01 | 3.53E-01 |
| IL-23 | 5.78E-01 | 7.51E-01 | 7.03E-01 | 8.31E-01 |
| IL-6 | 1.35E-01 | 1.96E-01 | 7.68E-01 | 8.32E-01 |
| CCL2 | 9.75E-01 | 9.75E-01 | **6.46E-03** | **2.80E-02** |
| IL-1β | **3.32E-02** | 5.39E-02 | 1.73E-01 | 2.81E-01 |
| IFN-β | **8.54E-03** | **1.59E-02** | **2.78E-02** | 9.04E-02 |
| IL-17A | **6.94E-03** | **1.50E-02** | 1.23E-01 | 2.29E-01 |
| IL-10 | 7.93E-01 | 8.59E-01 | 9.74E-01 | 9.74E-01 |
| TNF-α | **8.84E-04** | **3.83E-03** | **4.98E-02** | 1.08E-01 |
| CSF2 | 7.66E-01 | 8.59E-01 | 5.12E-01 | 6.65E-01 |
| IFN-γ | **3.03E-03** | **9.84E-03** | **4.91E-02** | 1.08E-01 |
| IL-12p70 | **7.96E-04** | **3.83E-03** | **5.20E-04** | **5.65E-03** |

\*Values are ordered according to expression level of measured cytokines in UPEC-infected prostates at 1 day PI. *p*-values were determined by Mann-Whitney nonparametric test, and corrected for multiple testing to determine *q*-values (false discovery rate adjusted *p*-value). *p*-values and *q*-values <0.05 are shown in bold.

**Table 8:** Statistical testing of analytes expressed during primary polymicrobial infection or challenge infection with UPEC or *E. faecalis*.

| *Analyte | Polymicrobial primary vs<br>UPEC challenge |  | Polymicrobial primary vs<br>*E. faecalis* challenge |  |
| --- | --- | --- | --- | --- |
|  | *p*-value | *q*-value | *p*-value | *q*-value |
| IL-1α | **6.64E-05** | **2.88E-04** | **2.10E-10** | **9.10E-10** |
| IL-27 | 1.52E-01 | 1.62E-01 | **3.04E-02** | **3.59E-02** |
| IL-23 | 8.79E-02 | 1.04E-01 | 9.73E-02 | 9.73E-02 |
| IL-6 | **2.71E-04** | **7.04E-04** | **4.44E-05** | **9.62E-05** |
| CCL2 | **2.33E-05** | **1.51E-04** | **4.38E-13** | **5.69E-12** |
| IL-1β | **3.96E-06** | **5.15E-05** | **2.21E-12** | **1.44E-11** |
| IFN-β | 1.62E-01 | 1.62E-01 | **4.24E-02** | **4.60E-02** |
| IL-17A | **2.10E-03** | **3.90E-03** | **3.32E-07** | **8.63E-07** |
| IL-10 | **2.13E-02** | **3.08E-02** | **1.38E-04** | **2.57E-04** |
| TNF-α | **1.23E-04** | **3.99E-04** | **2.87E-10** | **9.34E-10** |
| CSF2 | **8.35E-03** | **1.36E-02** | **2.44E-04** | **3.96E-04** |
| IFN-γ | **4.54E-02** | 5.90E-02 | **4.02E-04** | **5.80E-04** |
| IL-12p70 | **5.60E-04** | **1.21E-03** | **2.75E-03** | **3.58E-03** |

\*Values are ordered according to expression level of measured cytokines in UPEC-infected prostates at 1 day PI. *p*-values were determined by Mann-Whitney nonparametric test, and corrected for multiple testing to determine *q*-values (false discovery rate adjusted *p*-value). *p*-values and *q*-values <0.05 are shown in bold.

### Innate immune cells infiltrate UPEC-infected prostates more robustly

To assess immune cell infiltration induced by chemokine expression following bacterial infection, we instilled UPEC or *E. faecalis* into male mice and analyzed digested prostate tissue by flow cytometry at 24 hours PI (gating strategy in **Figure S3**). Following infection with either strain, the total number of cells in the prostate was unchanged compared to naïve animals (**Figure 4A,** compare orange to black dots). CD45^+^ immune cells were significantly higher only in UPEC-infected prostates, and not in *E. faecalis*-infected tissue, compared to naïve prostates (**Figure 4B**). In contrast to UPEC infection of the bladder (Zychlinsky Scharff et al., 2019), eosinophils only modestly infiltrated the prostate in primary UPEC infection and were not different from the naïve state in *E. faecalis* infection (**Figure 4C**). Similar to the female UTI model, UPEC infection induced robust neutrophil and monocyte-derived cell infiltration, which was significantly elevated above the numbers of these cells in naïve tissue (**Figure 4D**). Monocyte-derived cells were both MHC II^−^ and MHC II^+^, likely representing recently infiltrated cells (MHC II^−^) and those undergoing maturation to macrophages (MHC II^+^) (Mora-Bau et al., 2015). In *E. faecalis* infection, no populations were significantly elevated over naïve levels after correction for multiple testing. Finally, antigen-presenting cells, such as resident macrophages, CD11b^+^ dendritic cells (DCs), and CD103^+^ DCs were not different from naïve levels at 24 hours PI in either infection (**Figure 4E**).

**Figure 4.**
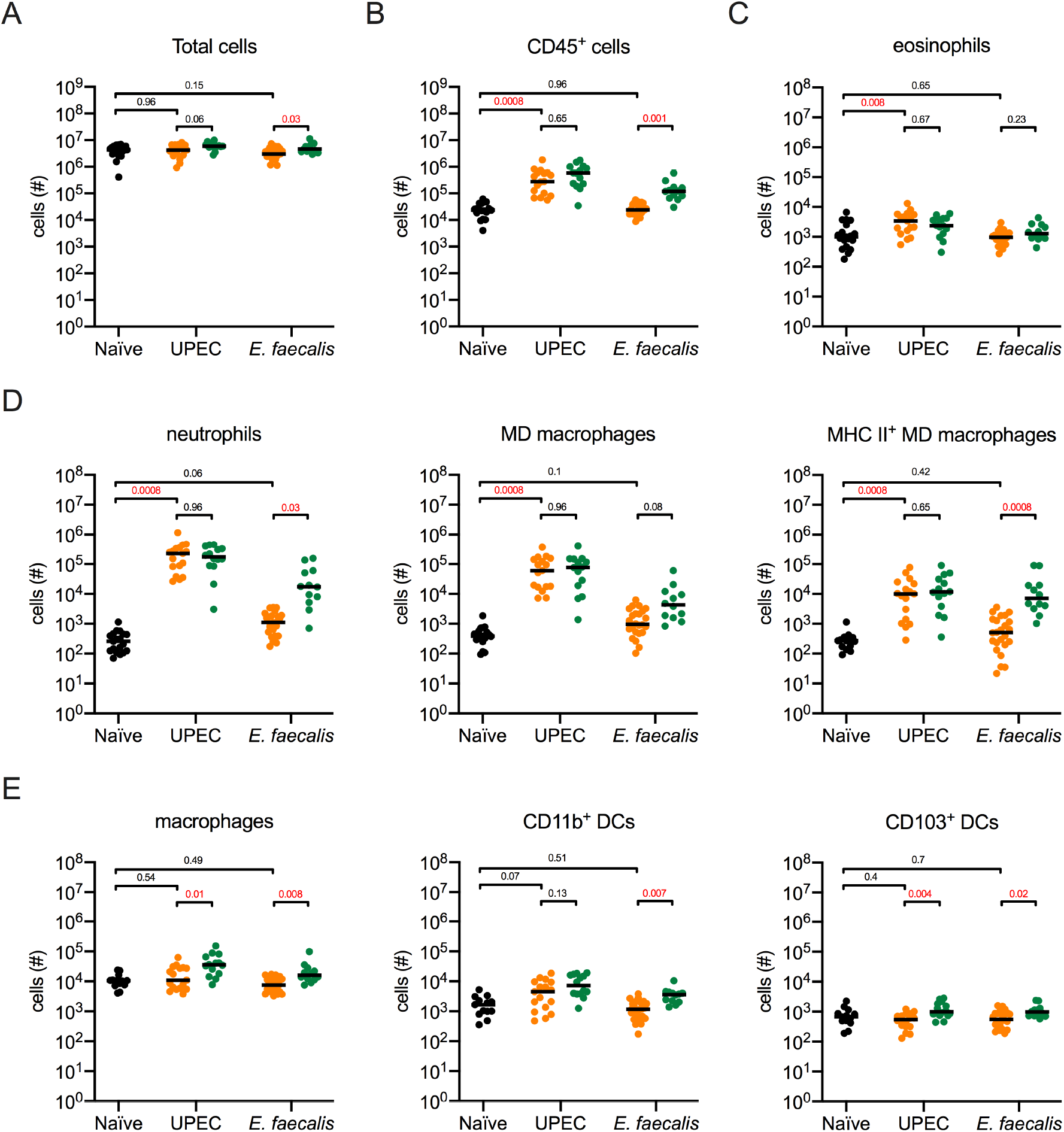
UPEC infection induces a greater immune cell infiltration in the prostate than *E. faecalis*. 6-8-week old male C57BL/6 mice were infected with 10^7^ CFU of either red fluorescent UPEC strain UTI89-RFP-kan^R^ or *E. faecalis* OG1RF stained with wheat germ agglutinin conjugated to Alexa Fluor 594. At 24 hours post-primary or post-challenge infection, mice were sacrificed, and prostates analyzed by flow cytometry. Naïve mice were included for baseline determination. **Figure S3** depicts the gating strategy used for these experiments. Graphs depict (**A**) total prostate cells, (**B**) total CD45^+^ immune cells, and (**C-E**) total specified immune cell populations. MD – monocyte-derived. Data are pooled from 2-3 experiments, with 5-7 mice per group in each experiment. Each dot represents one mouse, lines are medians, black dots depict naïve mice, orange dots denote values from primary infection, and green dots represent data from challenge infection. Statistical significance was determined using the non-parametric Mann-Whitney test for unpaired data and *p*-values were corrected for multiple testing using the false discovery rate (FDR) method. *q*-values meeting the criteria for statistical significance (*q* <0.05) are depicted in red.

Given that cytokine responses were similar or elevated in response to challenge infection in monomicrobial infections, we measured immune cell infiltration into the prostate 24 hours post-challenge infection. We infected mice with 10^7^ CFU of UPEC or OG1RF and monitored infection via urine collection. One month after infection, following antibiotic treatment and a washout period, mice with sterile urine were challenged with isogenic strains of UPEC or *E. faecalis*. Prostates were collected, digested, and stained for flow cytometric analysis 24 hours PI, as above. Whereas the total numbers of prostate cells and total CD45^+^ cells were not different between UPEC primary and challenge infection, *E. faecalis* challenged prostates had more total cells and greater CD45^+^ cell infiltration compared to the primary infection (**Figure 4A-B,** compare orange to green dots for each bacterial strain). Eosinophil infiltration was not different in comparison to primary infection with either strain, and given how little this population changed overall, these cells may not play an important role in the response to prostate infection (**Figure 4C)**. In UPEC infection, although neutrophils and both subsets of monocyte-derived macrophages were elevated in a primary infection compared to naïve levels, these populations were not further increased in UPEC challenge compared to the primary infection (**Figure 4D**). Conversely, neutrophil and MHC II^+^ monocyte-derived macrophage infiltration was significantly elevated in *E. faecalis*-challenged animals compared to the numbers observed 24 hour post-primary infection (**Figure 4D**). Resident macrophages and CD103^+^ DC populations increased in mice challenged with UPEC or *E. faecalis*, whereas CD11b^+^ DCs increased only in *E. faecalis*-challenged mice (**Figure 4E**). Overall, these results demonstrate that myeloid cell recruitment is robust in response to a first UPEC infection but does not change in the context of a challenge or recurrent prostatitis. By contrast, a primary *E. faecalis* infection is largely silent and challenge infection by this Gram-positive organism is required for robust myeloid cell infiltration.

### The phagocyte compartment engulfs more UPEC than E. faecalis early in infection

Dependent upon the infecting organism, significant innate immune cell infiltration was evident post-primary or challenge infection. We next determined whether this impacted bacterial phagocytosis during primary or challenge infection. In infected organs, UTI89-RFP-kan^R^ is easily detectable by flow cytometry (Mora-Bau et al., 2015; Zychlinsky Scharff et al., 2019), however we did not have a red fluorescent strain of *E. faecalis*. In testing a GFP-expressing *E. faecalis* strain, we encountered challenges similar to those with using GFP-expressing UPEC in the bladder, in that tissue autofluorescence masked the GFP signal (**Figure S4**). Therefore, we stained *E. faecalis* bacterial cell walls with wheat germ agglutinin conjugated to Alexa-Fluor 594 to visualize immune cells that had taken up *E. faecalis*. Overall, in primary and challenge infection, we observed greater numbers of UPEC-infected immune cells in the prostate in comparison to the number of *E. faecalis*-infected cells (**Figure 5A-C**). When we considered bacterial uptake between primary and challenge infection, we observed that there were no differences in the number of UPEC^+^ cells in any of the populations we measured (**Figure 5A-C**). However, in *E. faecalis* infection, we observed that the total number of CD45^+^ cells containing bacteria was increased in the challenge infection, compared to primary infection (**Figure 5A**). Among specific phagocyte populations, significant increases in the number of resident macrophages and mature (MHC II^+^) monocyte-derived cells containing *E. faecalis* were observed 24 hours post-challenge infection (**Figure 5B-C**).

**Figure 5.**
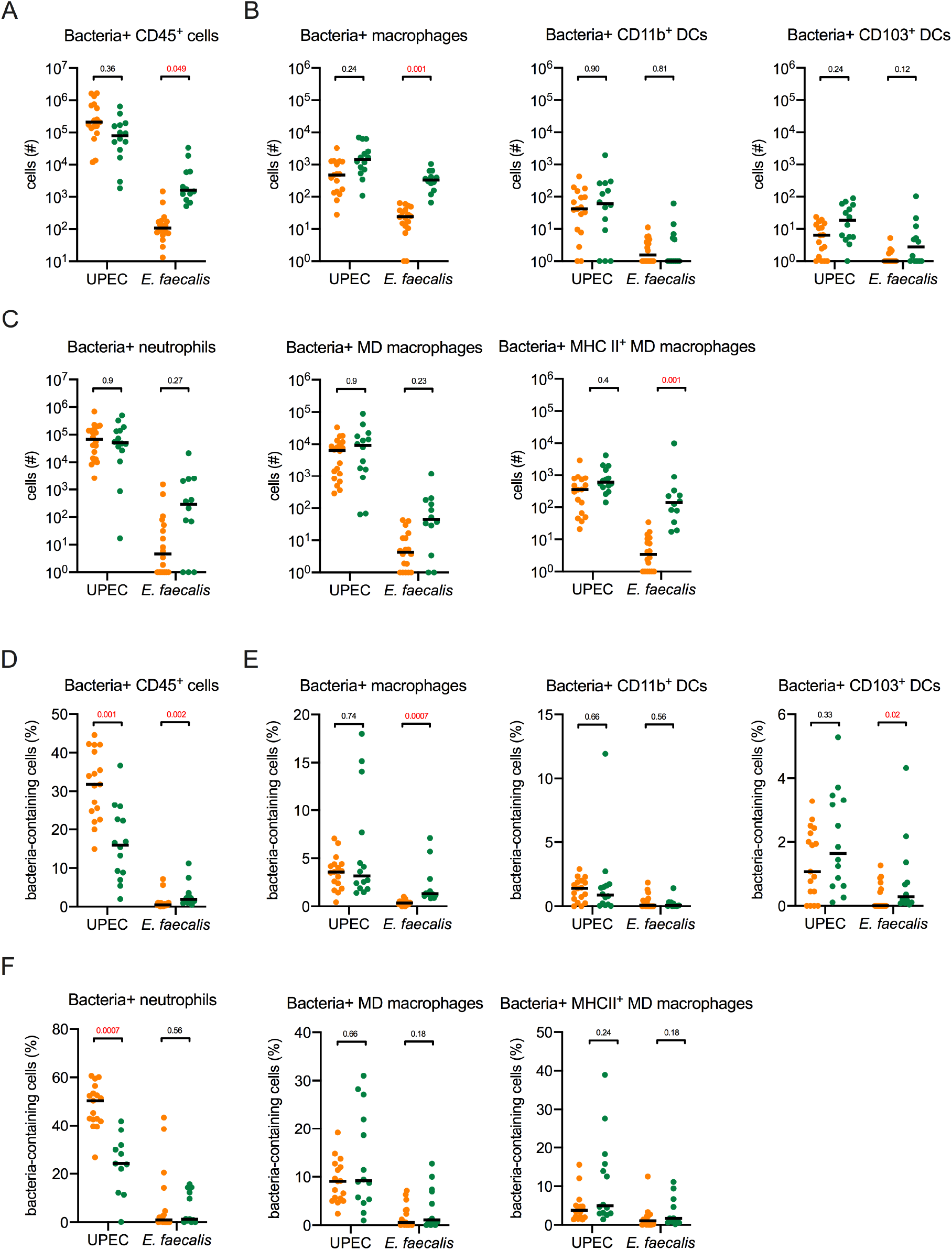
Phagocytosis is altered by repeated infection. 6-8-week old male C57BL/6 mice were infected with 10^7^ CFU of either red fluorescent UPEC strain UTI89-RFP-kan^R^ or *E. faecalis* OG1RF stained with wheat germ agglutinin conjugated to Alexa Fluor 594. At 24 hours post-primary or post-challenge infection, mice were sacrificed, and prostates analyzed by flow cytometry. Graphs show the (**A-C**) absolute number or (**D-F**) percentage of cells positive for red-fluorescent bacteria within the specified immune cell populations depicted. MD – monocyte-derived. **Figure S3** shows the gating strategy used for these experiments. Data are pooled from 2-3 experiments, with 5-7 mice per group in each experiment. Each dot represents one mouse, lines are medians, black dots depict naïve mice, orange dots denote values from primary infection, and green dots represent data from challenge infection. Statistical significance was determined using the non-parametric Mann-Whitney test for unpaired data and *p*-values were corrected for multiple testing using the false discovery rate (FDR) method. *q*-values meeting the criteria for statistical significance (*q* <0.05) are depicted in red.

We then considered whether phagocytic capacity was altered between primary and challenge infection by calculating the percentage of cells containing bacteria for the indicated immune cell populations in the prostate. We observed that the percentage of CD45^+^ immune cells containing bacteria was decreased in UPEC challenged-mice, whereas it was increased in *E. faecalis*-challenged mice (**Figure 5D)**. A greater proportion of resident immune cell populations, such as macrophages and CD103^+^ DCs contained *E. faecalis* following challenge infection (**Figure 5E**) While these resident populations were not different in UPEC-challenged mice compared to primary infection, a significant decrease in the percentage of neutrophils containing bacteria was evident, suggesting that the capacity of neutrophils to take up bacteria post-challenge is impaired in UPEC infection (**Figure 5F**).

### Effector cells efficiently infiltrate the prostate during challenge infection

We next determined whether the differences we observed in cytokine expression and innate immune cell numbers impacted effector cell infiltration during UPEC or *E. faecalis* primary and challenge infection. Using flow cytometry, we analyzed the numbers of T cells, NK cells, NK T cell, and innate lymphoid cells (ILCs) present in prostates of naïve, primary, and challenge-infected mice. In a primary infection, T cells, comprised almost entirely of CD4^+^ T cells, were significantly elevated over naïve levels in the prostate following UPEC, but not *E. faecalis* infection (**Figure 6A-B,** compare orange to black dots). The global CD3^+^ T cell population, and specifically the CD4^+^ T cell pool, was significantly increased upon challenge with either bacterial strain over their respective primary infection levels, suggesting that a memory response was induced (**Figure 6A-B,** compare green to orange dots). NK cells were also increased in challenge infection compared to primary infection with either organism (**Figure 6C**). NK T cells were significantly elevated in prostates following primary UPEC infection and *E. faecalis* challenge (**Figure 6C**). Finally, the total number of innate lymphocytes was unchanged at all timepoints, suggesting that these cells do not play a role in prostatitis as no infiltration was induced (**Figure 6C**).

**Figure 6.**
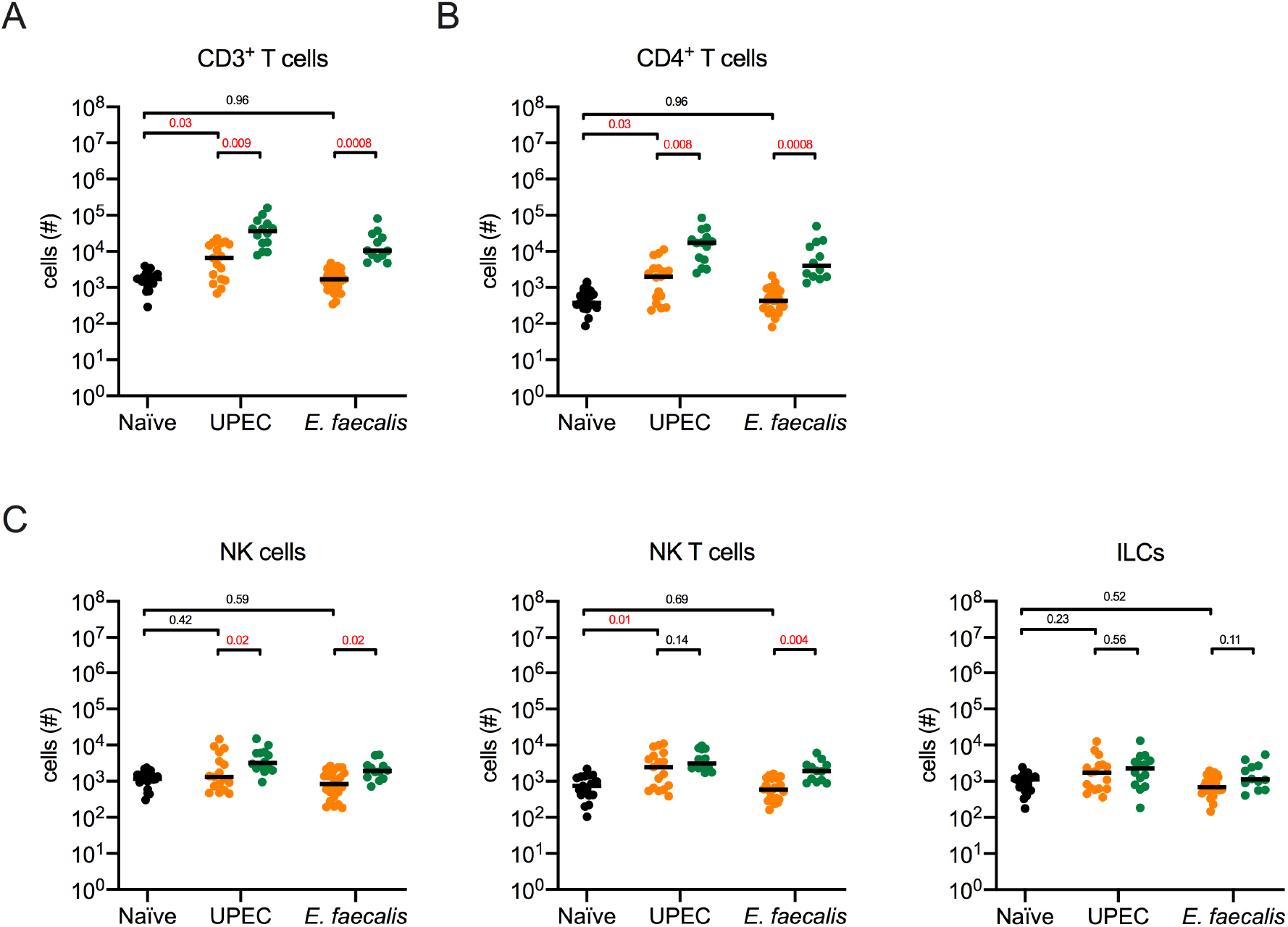
Effector immune cell infiltration is increased in prostates challenged with UPEC or *E. faecalis*. 6-8-week old male C57BL/6 mice were infected with 10^7^ CFU of either red fluorescent UPEC strain UTI89-RFP-kan^R^ or *E. faecalis* OG1RF stained with wheat germ agglutinin conjugated to Alexa Fluor 594. At 24 hours post-primary or post-challenge infection, mice were sacrificed, and prostates analyzed by flow cytometry. Naïve mice were included for baseline determination. **Figure S3** depicts the gating strategy used for these experiments. Graphs depict (**A**) CD3^+^ T cells, (**B**) CD3^+^ CD4^+^ T cells, and (**C**) total specified immune cell populations. Data are pooled from 2-3 experiments, with 3-7 mice per group in each experiment. Each dot represents one mouse, lines are medians, black dots depict naïve mice, orange dots denote values from primary infection, and green dots represent data from challenge infection Statistical significance was determined using the non-parametric Mann-Whitney test for unpaired data and *p*-values were corrected for multiple testing using the false discovery rate (FDR) method. *q*-values meeting the criteria for statistical significance (*q* <0.05) are depicted in red.

### Prostate infection does not induce a protective immune response to bacterial challenge

Given the pronounced effector cell accumulation we observed during UPEC and *E. faecalis* infection, we hypothesized that this strong adaptive immune response would mediate protection against subsequent bacterial colonization. Indeed, a protective immune response is induced during female and male bladder infection with UPEC, although this response is not sterilizing and animals challenged with UPEC are still colonized but at significantly lower levels compared to a primary infection (Mora-Bau et al., 2015; Zychlinsky Scharff et al., 2019). Given the robust innate response in the prostate to UPEC infection compared to the near lack of response in *E. faecalis* infection, we investigated whether a protective adaptive immune response arises following infection with either uropathogen. We infected cohorts of male C57BL/6 mice with 10^7^ CFU UPEC, OG1RF, or with a 1:1 ratio of both strains. We determined the bacterial burden in the prostate and bladder 24 hours PI in a subset of animals after a primary infection, and monitored bacterial clearance over the course of one month in the remaining mice. Mice that received UPEC or *E. faecalis* were challenged with an isogenic strain of the same bacteria used for primary infection. Animals receiving a primary polymicrobial infection were challenged with either UPEC or *E. faecalis* to evaluate the impact of primary co-infection on development of an adaptive immune response to either bacterial strain. In the bladder, 24 hours post-challenge infection, the bacterial burden was significantly decreased compared to that in primary UPEC-infected bladders, as we previously reported (**Figure 7A**) (Zychlinsky Scharff et al., 2019). Surprisingly, prostate UPEC burdens, from the same mice, were nearly 10 times higher in challenged organs compared to the bacterial burden observed in prostates following the primary infection (**Figure 7A**). Bacterial CFU in prostates challenged with *E. faecalis* were similar to primary infection levels, while, quite surprisingly, the bladders from these same animals harbored nearly 100-fold greater bacterial burden than in primary infection (**Figure 7B**). In the polymicrobial infection scenario, quite strikingly, protection against UPEC was lost in the bladders of these mice (**Figure 7C**). In addition, protection against UPEC or *E. faecalis* was not observed in the prostates of mice that experienced a primary polymicrobial infection (**Figure 7C-D**). Together, these data support that *E. faecalis* plays a role in the suppression of adaptive immune responses, particularly in the bladder. Indeed, we never observed a reduction in bacterial burden in prostates after challenge infection in any scenario, supporting that although a non-sterilizing adaptive immune response is mounted in the bladder, specifically to UPEC, no protective adaptive response to infection develops following prostate colonization.

**Figure 7.**
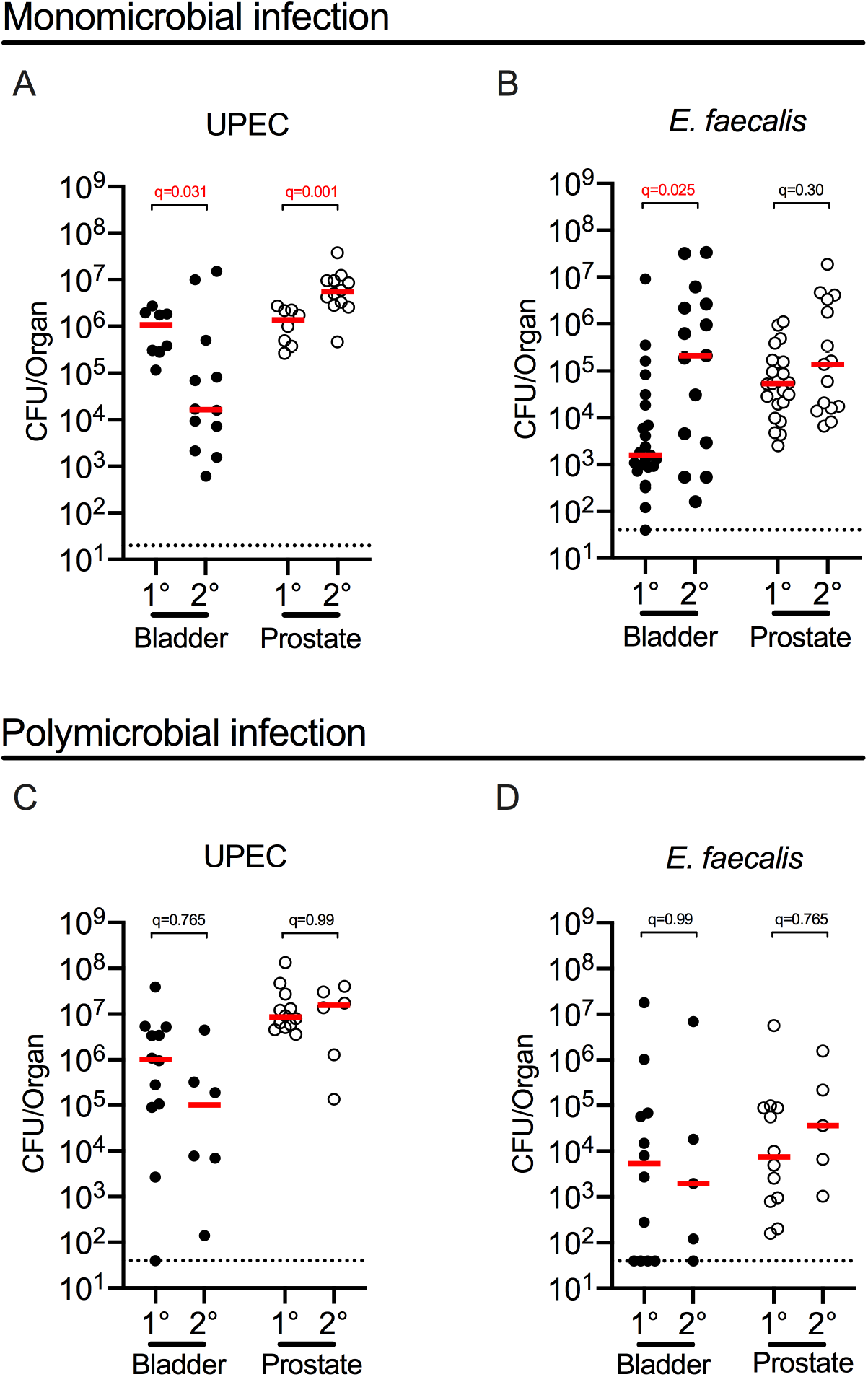
The prostate fails to mount a protective immune response against bacterial reinfection. 6-8-week old male C57BL/6 mice were infected with 10^7^ CFU of (**A**) UPEC strain UTI89-RFP-kan^R^, (**B**) *E. faecalis* strain OG1RF, or (**C**) a total of 10^7^ CFU of both bacteria strains in an approximate 1:1 ratio. One half of each group was sacrificed at 24 hours post-primary (1°) infection. At 25-30 days post-infection (PI), all remaining animals were treated with one or two cycles of antibiotics as described in the Materials and methods for 5 days, followed by a 3-5 day washout period. Mice with sterile urine were challenged with 10^7^ CFU of (**A**) UPEC strain UTI89-GFP-amp^R^, (**B**) *E. faecalis* strain OG1RF_intergenicRS00490RS00495::Tn, or in primary polmicrobially-infected mice, half of the cohort was challenged with (**C**) UTI89-GFP-amp^R^ and the other half was challenged (**D**) with OG1RF_intergenicRS00490RS00495::Tn. Mice were sacrificed 24 hours post-challenge infection (2°). Graphs depict CFU/organ. Each circle represents one mouse, filled circles represent bladder CFU, open circles represent prostate CFU. Red lines indicate the median value of each group and black dotted lines represent the limit of detection of the assay, 20 or 40 CFU/organ. Data are pooled from 2-4 experiments, each with 2-6 mice per group. CFU from 1° and 2° infection in bladder or prostate were compared using the non-parametric Mann-Whitney test for unpaired data. Within each infection scenario (UPEC, *E. faecalis*, polymicrobial) *p*-values were corrected for multiple testing using the false discovery rate (FDR) method. *q*-values meeting the criteria for statistical significance (*q* <0.05) are depicted in red.

## Discussion

One percent of all men will experience bacterial prostatitis in their lifetime and half of them will have chronic relapses in the months following infection. The only treatment option is antibiotics with regimens that may last from a few weeks to several months, increasing the risk of development of antibiotic resistance. Although the symptomatology and microbiological causes of bacterial prostatitis are reasonably well investigated, little is known about the prostate immune compartment and its role in protecting against infection. Here, we used a mouse model of male UTI to study the immune response in the prostate following catheter-mediated transurethral instillation of UPEC or *E. faecalis*. Mouse prostates were colonized for at least four weeks by UPEC or *E. faecalis*, despite an acute proinflammatory response, including robust neutrophil and monocyte-derived cell infiltration, in the case of UPEC infection. Notably, the immune response was the same or greater upon challenge infection with both strains, although surprisingly the prostate did not mount a protective immune response to challenge infection, regardless of a pronounced lymphocytic infiltration.

Both UPEC and *E. faecalis* displayed tropism for the prostate over the bladder at 24 hours PI, but *E. faecalis* persisted at higher levels in the prostate compared to the bladder up to two weeks PI. Despite persistent infection, *E. faecalis* did not elicit as strong an inflammatory response compared to UPEC. Differences in initial bacterial burden between UPEC and *E. faecalis* may be only partially responsible for the divergent cytokine response to the two organisms as *E. faecalis* can suppress NF-kB-mediated proinflammatory responses in myeloid cells, dampening proinflammatory cytokine pathways (Tien et al., 2017). Additionally, *E. faecalis*-mediated immunosuppression may promote colonization or fitness of other bacteria (Kao and Kline, 2019; Tay et al., 2016).

The greater cytokine response in the prostate over the bladder in the first two days PI was surprising, as we recently showed that cytokine levels are globally suppressed in the bladder following UPEC infection in male mice compared to female mice (Zychlinsky Scharff et al., 2019). These findings suggest that during infection, cytokine expression in the prostate remains localized and mechanisms that result in inadequate cytokine expression in the bladder are restricted to this tissue. Additionally, we observed different baseline levels of IL-6 and IL-1β between prostate and bladder, which may lead to divergent innate immune responses as these tissues may be differently primed to respond to infection. However, insights into the prostate-specific role of IL-6 and IL-1β are needed to verify this hypothesis. In fact, these cytokines underpin early stage inflammatory processes, such as neutrophil recruitment, and adaptive immune processes, including T_h_17 polarization (Bettelli et al., 2006; Netea et al., 2010; Rincon, 2012). Albeit often synergistically correlated, IL-6 and IL-1β may also function independently (McGeough et al., 2012). As an added layer of complexity, the inflammatory role of IL-6 is diverse. For example, IL-6^−/−^ mice infected with *E. coli*, *C. albicans,* or *M. tuberculosis* have significantly reduced neutrophil numbers and survival, with a poorer inflammatory response (Dalrymple et al., 1996; Ladel et al., 1997; Romani et al., 1996). However, aerosol exposure or intraperitoneal injection of endotoxin into IL-6^−/−^ mice induces local and systemic increases in TNF-α, CXCL2, and neutrophil infiltration in the lungs at significantly higher levels than in wildtype mice, in which IL-6 production is observed (Xing et al., 1998). Therefore, the role of IL-6 in immunity of the prostate may be quite complex. Furthermore, despite differences in the magnitude of cytokine response between UPEC and *E. faecalis* infection, IL-1α, IL-6, CCL2, and IL-1β were significantly increased in the prostate over the bladder for two days PI with either bacteria, suggesting that common antibacterial innate immune pathways may be elicited in the prostate in which these cytokines are equally pivotal.

Compared to the bladder (Zychlinsky Scharff et al., 2019), the acute immune response to UPEC infection in the prostate was composed almost entirely of neutrophils and monocytes. The greater neutrophil involvement may be due to the copious secretion of IL-1α in the prostate, which recruits neutrophils in other tissues (Chen et al., 2007). IL-1α was five times higher than any other cytokine in UPEC infection, and notably absent in *E. faecalis* infection. The role of IL-1α in immunity is less clear than IL-1β, even though they both signal through interleukin 1 receptor-MyD88-dependent inflammatory pathways (Dinarello, 2009). Constitutively active and normally sequestered in the cell cytoplasm, IL-1α is translocated to the plasma membrane during stress conditions and released by necrotic cells, inducing tissue inflammation (Garlanda et al., 2013). It is rapidly inducible in cells of hematopoietic origin, such as macrophages or dendritic cells, and expressed by stromal cells such as keratinocytes or endothelial cells, as well (Di Paolo and Shayakhmetov, 2016). IL-1α may also induce local inflammation in an “alarmin”-like fashion. Notably, a sterile inflammatory response to necrotic HMGB1-deficient cells leads to IL-1α-dependent acute neutrophil infiltration in two different *in vivo* models (Chen et al., 2007). Similar mechanisms may be at play in UPEC-infected prostates, in which the release of IL-1α from necrotic cells and tissue damage provoked by UPEC invasion mediates immune cell infiltration. Thus, IL-1α-activated stromal cells would secrete proinflammatory cytokines in an autocrine inflammatory loop to enhance neutrophil and monocyte recruitment via the release of chemotactic mediators. Of great clinical interest, the use of anakinra, an IL-1R antagonist, greatly reduced infection-associated pathology in a mouse model of cystitis (Ambite et al., 2016), supporting that mechanisms targeting the immune response may provide relief to patients suffering from prostatitis.

We observed a pronounced lack of protective immunity in prostates of UPEC- and *E. faecalis* infected mice, despite that protection was observed in the bladder of the same mice following UPEC infection. This suggests that lymphocytes in the two organs respond differently to challenge infection due to the divergent cytokine environment, that lymphocytes specific for bacterial antigens do not develop or infiltrate, or that bacteria in the prostate are simply inaccessible to responding immune cells. Previously, we showed that the bladder adaptive immune response upon challenge infection is significantly improved when macrophages are depleted prior to primary infection (Mora-Bau et al., 2015). This subversive role of macrophages may also occur in the prostate. For example, in UPEC or *E. faecalis* infection, a vast majority of bacteria engulfing cells were resident or monocyte-derived macrophages, and few bacteria could be found in tissue DCs.

In summary, we found that mouse prostates remained chronically infected for at least four weeks by UPEC or *E. faecalis*. While infection induced strong soluble or cellular innate responses, ultimately the prostate was not protected against challenge infection with either bacteria. This finding may underlie the high rate of chronic prostatitis observed in men. Overall, these results support that the prostate is responsive to bacterial colonization but fails to develop lasting protection in the face of recurrent infection. Understanding mechanisms underlying this ineffectual response may lead to the identification of targetable pathways for immunotherapeutic approaches in prostatitis.

## Materials and methods

### Study approval

This study was conducted using a preclinical mouse model in controlled laboratory experiments to test the hypothesis that the immune response to infection with *E. coli* or *E. faecalis* is similar between the bladder and prostate. Animals were assigned to groups by random partition into cages. In each experiment, a minimum of three and a maximum of six animals constituted an experimental group and all experiments were repeated two to ten times. Data were pooled before statistical analysis. We have observed that abnormal kidneys negatively impact resolution of infection; thus, in all of our studies, we have established *a priori* that mice with atrophied, enlarged, or markedly pale kidneys at the time of sacrifice are to be excluded from all analyses. Endpoints were determined before beginning experiments and researchers were not blinded to experimental groups.

### Ethics Statement and mice

Animal experiments were conducted in accordance with the protocol number 2016-0010, approved by the *Comités d’Ethique pour l’Expérimentation Animale* at Institut Pasteur (the ethics committee for animal experimentation), in observation of the European Directive 2010/63 EU. In this study, C57BL/6 male mice between the ages of 6-12 weeks from Charles River Laboratories France were used. Mice were anaesthetized by intraperitoneal injection of 100 mg/kg ketamine and 5 mg/kg xylazine and sacrificed by carbon dioxide inhalation.

### Urinary tract infection

Male mice were anesthetized as above, catheterized transurethrally, and infected with a total of 10^7^ colony forming units (CFU) of single or mixed bacterial inocula of indicated strains of uropathogenic *E. coli* (UTI89-GFP-amp^R^ or UTI89-RFP-kan^R^) and/or *E. faecalis* (OG1RF, OG1RF-GFP, OG1RF_intergenicRS00490RS00495::Tn) in 50 µL PBS as previously described (Dale et al., 2018; Debroy et al., 2012; Dunny et al., 1978; Hung et al., 2009; Kristich et al., 2008; Mora-Bau et al., 2015; Zychlinsky Scharff et al., 2017). Antibiotic resistance and concentrations used are shown in **Supplementary Table 1**.

Urine was collected 24 hours PI and every 2-5 days thereafter. Two µL of urine were diluted into 8 µL PBS spotted on agar plates containing antibiotics as appropriate. The qualitative presence of any bacterial growth was counted as positive for infection. The limit of detection (LOD) for this assay is 500 bacteria per mL of urine. At 25-30 days PI, we treated mice with antibiotics, dependent upon the initial infection. UPEC-infected mice were given 8 mg/mL trimethoprim-sulfamethoxazole (TMP-SMX, Avemix) in the drinking water. Mice infected with *E. faecalis* were intraperitoneally injected with 10 mg of carbenicillin in 100 µL PBS (Baker and Androle, 1973; Libke et al., 1973). Mice co-infected with both organisms received both antibiotics. Antibiotic treatments were administered for five days and were followed by a washout period of 3-6 days prior to challenge infection. In the case that a majority of animals remained infected after the first washout period, a second round of antibiotics was administered. Following the washout period, monomicrobial-infected mice were challenged with isogenic strains of the bacteria used for primary infection. Polymicrobial-infected mice were challenged with either UPEC or *E. faecalis* isogenic strains.

### Determination of bacterial burden and cytokine analysis

To determine CFU in infected organs, mice were sacrificed 1, 2, 14, or 42 days PI and bladders and prostates were removed. Organs were immediately placed in 1 mL of cold PBS, then homogenized with a PreCellys® 24 homogenizer. 100 µL from homogenized organs were serially diluted and plated on appropriate complete medium (LB agar for UPEC and BHI for *E. faecalis*) with antibiotics as appropriate. The LOD for organ CFU is 20 or 40 bacteria per organ, depending upon the number and volume of dilutions plated, and is indicated by a dotted line in graphs. All sterile organs are reported at the LOD.

Homogenates were clarified by microcentrifugation (17,000xg; 4°C; 5 minutes) and supernatants were stored at −20°C until analysis for cytokine levels in low protein binding plates. After thawing on ice, samples were centrifuged again to remove remaining cell debris prior to analysis (200g; 4°C; 5 minutes). To avoid inter-assay variability, whenever possible, samples were analyzed at the same time with the LEGENDplex^TM^ Mouse Inflammation Panel (Biolegend, USA). The assay was performed following manufacturer’s instructions for measuring serum or plasma samples using a V-bottom plate, with one modification: the first incubation on the shaker was extended, from 2 hours at room temperature to overnight at 4°C. Samples were acquired on an LSRFortessa^TM^ (BD Biosciences, USA) and analyzed using version 7.1 of the LEGENDplex^TM^ Data Analysis Software (Biolegend, USA) for Mac OS X.

### Flow cytometry

Mice were sacrificed 24 hours post-primary or post-challenge infection and the entire prostate removed. Single cell suspensions were prepared following modification of a protocol for preparation of bladder single cell suspensions previously described (Zychlinsky Scharff et al., 2019). Briefly, minced prostates were incubated in 0.34 Units/mL Liberase TM (Roche, France), diluted in PBS at 37°C for 45 minutes, with manual agitation every 15 minutes. Digested tissue was passed through a 100 µm filter (Miltenyi Biotec, Germany), washed, blocked with FcBlock (BD Biosciences, USA), and immunostained with the indicated antibodies (**Supplementary Table 2**). Samples were acquired on an LSRFortessa^TM^ (BD Biosciences, USA) and analyzed using FlowJo Version 10 (FlowJo, USA) for Mac OS X. Prior to cytometer acquisition, 10 µL of stained sample were added to 20 µL AccuCheck Counting Beads (Invitrogen) in 200 µL PBS to measure total cell counts in the prostate. The gating strategies used are depicted in **Supplementary Figure 3**.

### Fluorescent staining of E. faecalis

To detect *E. faecalis* by flow cytometry, bacteria were stained with wheat germ agglutinin conjugated to Alexa Fluor 594 (Thermo Fisher, France) just prior to instillation. Wheat germ agglutinin conjugate stock solution was prepared as per manufacturer’s instructions in PBS without addition of sodium azide, aliquoted, and frozen at −20°C. Prior to staining, the stock solution was thawed, centrifuged for 10 seconds at 17,000x*g* to eliminate protein aggregates and the supernatant was used for staining. Wheat germ agglutinin conjugate was mixed with the bacterial inoculum at a working concentration of 10 µg/mL and incubated statically for 10 minutes at 37°C. The inoculum was spun at 17,000x*g* for one minute, the supernatant removed, and pellet resuspended in PBS for a total of three washes. The bacterial concentration was adjusted to 2×10^8^ CFU/mL in PBS for infection.

### Statistical analysis

Statistical analysis was performed in GraphPad Prism 8 (GraphPad, USA) for Mac OS X applying the non-parametric Wilcoxon test for paired data, the non-parametric Mann-Whitney test for unpaired data. When more than 2 groups were being compared or to correct for comparisons made within an analysis or experiment, calculated *p*-values were corrected for multiple testing with the false discovery rate (FDR) method (https://jboussier.shinyapps.io/MultipleTesting/), to determine the *q*-value (false discovery rate adjusted *p*-value). All calculated *q*-values are shown in the figures, and those that met the criteria for statistical significance (*q* <0.05) are denoted with red text. Qlucore Omics Explorer 3.6 software (Qlucore, Sweden) was used to calculate *q*-values of cytokine protein expression using the non-parametric Mann-Whitney with correction for multiple testing. Calculated *p*-values and *q*-values are shown in **Tables 1-8**, in which bold text denotes *p*-values or *q*-values <0.05

## Author contributions

Conceptualization: FL, MAI; Methodology: FL, MR, TC, MAI; Investigation and data analysis: FL, MR, TC, MAI; Writing - Original Draft: FL, MAI; Writing - Review & Editing: FL, MR, TC, MAI; Funding Acquisition: MAI; Supervision: MAI.

## Acknowledgments

We gratefully acknowledge insightful discussions and support from Livia Lacerda Mariano, Camila Rosat Consiglio, Jérémy Boussier, Kao Hsien-Neng, Choo Pei Yi, Giovanna Barba, Gerald Spaeth, and Maria Luisa Pisanelli. We thank Dr Kimberly Kline for sharing *E. faecalis* strains and critical feedback on the project and manuscript. Drs Karen Sfanos and Darragh Duffy provided much appreciated constructive critiques on the manuscript, as well. We also thank the University of Glasgow, including Drs Simon Milling and Robert Nibbs, for the academic opportunity offered through the integrated Masters (MSci) program with work placement in Life Sciences. Funding: This study was supported by funding from the *Agence Nationale de la Recherché* (French National Research Agency) ANR-17-CE17-0014.

## Supplemental Figures and Legends

**Supplementary Table 1:** Antibiotic concentrations used in this study.

| Strain name | Antibiotic resistance, agar plate concentration |
| --- | --- |
| UTI89-GFP-amp<sup>R</sup> | Ampicillin, 100 μg/mL |
| UTI89-RFP-kan<sup>R</sup> | Kanamycin, 50 μg/mL |
| *E. faecalis* strain OG1RF | rifampicin, 250 μg/mL |
| OG1RF-GFP | rifampicin, 250 μg/mL |
| OG1RF_intergenicRS00490RS00495::Tn | rifampicin, 250 μg/mL, chloramphenicol 7 μg/mL |

**Supplementary Table 2:** Antibodies used in this study.

| Molecule | Clone | Vendor |
| --- | --- | --- |
| B220 (CD45R) | RA3-6B2 | BD Biosciences |
| CD11b | M1/70 | BD Biosciences |
| CD3 | 145-2C11 | BD Biosciences |
| CD4 | RM4-5, GK1.5 | BD Biosciences |
| CD45 | 30-F11 | BD Biosciences |
| CD64 | X54-5/7.1 | BD Biosciences |
| CD90.2 | 53.2.1 | eBioscience |
| CD103 | M290 | BD Biosciences |
| F4/80 | CI:A3-1 | AbD Serotec |
| Ly6C/Ly6G (Gr-1) | RB6-8C5 | BD Biosciences |
| MHC-II (I-A/I-E) | M5 or 114.15.2 | eBioscience |
| NK1.1 | PK136 | BD Biosciences |
| SiglecF | E50-2440 | BD Biosciences |

**Figure S1.**
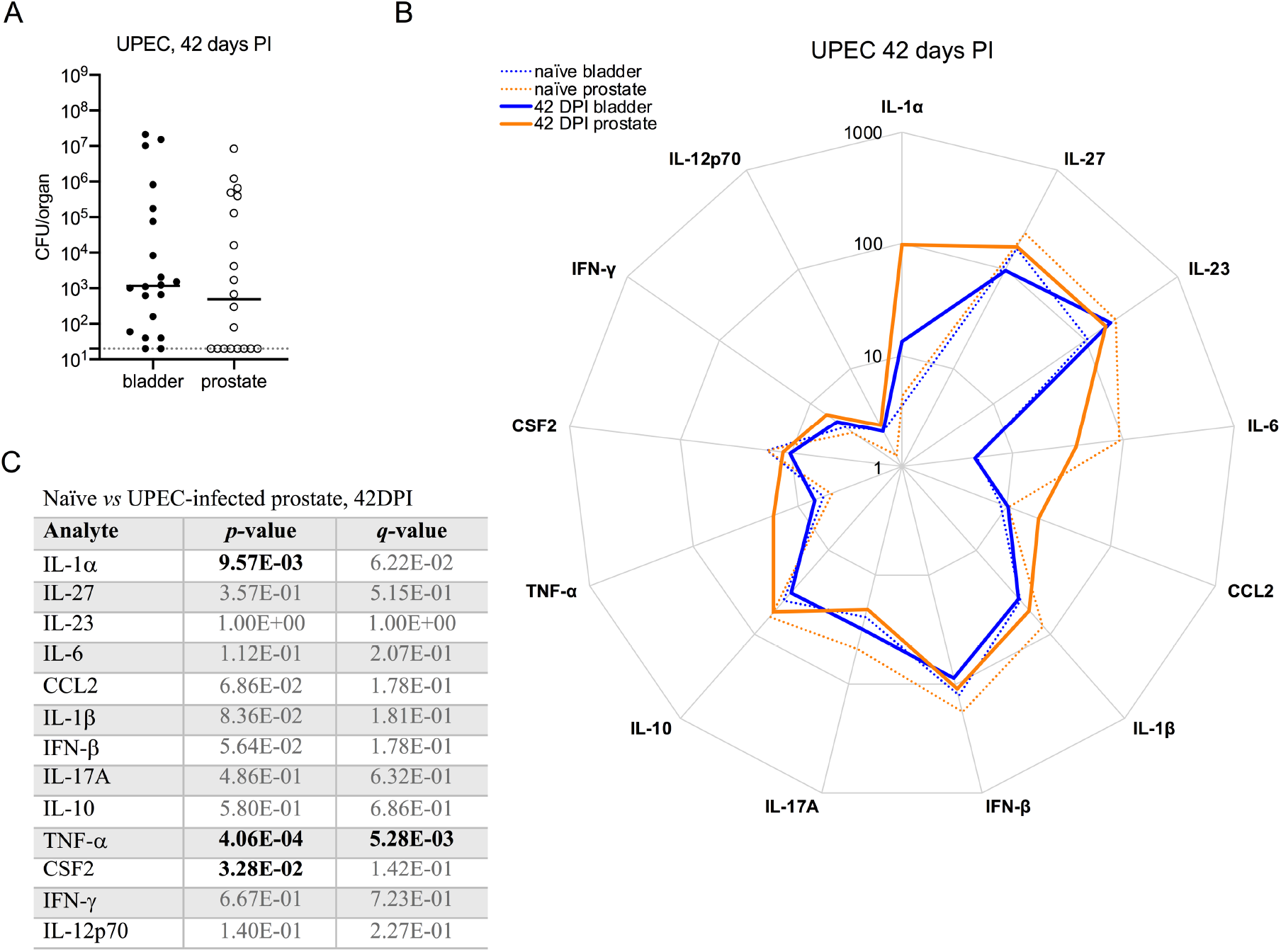
UPEC persists in the bladder and prostate up to 6 weeks. 6-8-week old male C57BL/6 mice were infected with 10^7^ CFU of UPEC strain UTI89-RFP-kan^R^ and sacrificed 42 days post-infection (PI). (**A**) The graph depicts CFU/organ in homogenized bladder and prostate tissue 42 days PI. The dotted line denotes the limit of detection for the assay. (**B**) 13 cytokines were measured in the supernatant of homogenized bladders and prostates by multiplex analysis. The spider plot shows the median values of absolute cytokine levels (pg/mL) on a log-scale in infected animals at 42 days PI (solid lines) together with naïve levels (dotted lines) for comparison to baseline. All data are plotted from 3 experiments, each with 5-7 mice Statistical significance was determined using the non-parametric Mann-Whitney test for unpaired data and *p*-values were corrected for multiple testing using the false discovery rate (FDR) method. (**C**) The table lists calculated *p*- and *q*-values for each analyte.

**Figure S2.**
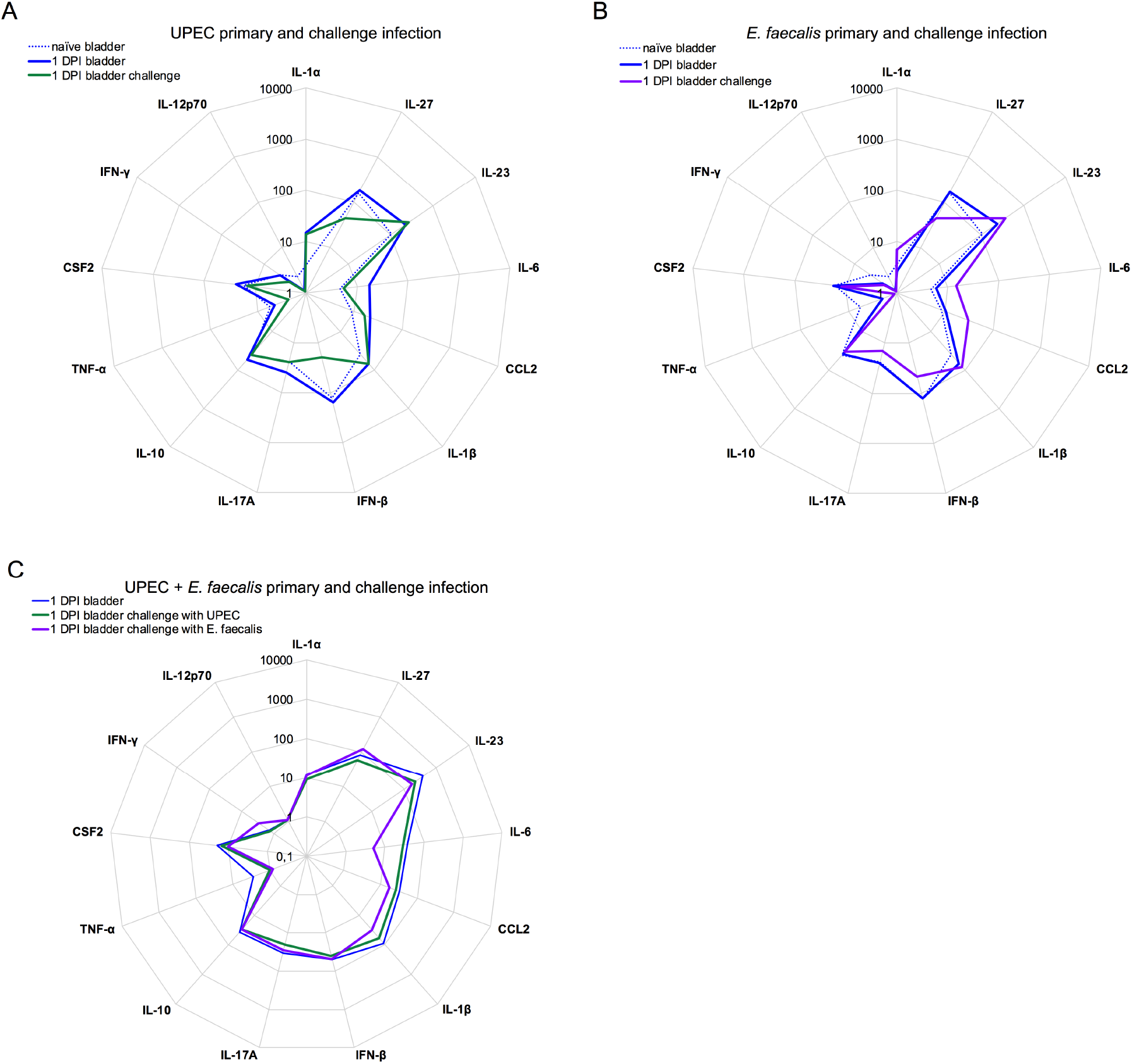
The cytokine response is altered following challenge infection. 6-8-week old male C57BL/6 mice were infected with 10^7^ CFU of (**A**) UPEC strain UTI89-RFP-kan^R^, (**B**) *E. faecalis* strain OG1RF, or (**C**) a total of 10^7^ CFU of both bacteria strains in an approximate 1:1 ratio. At 25-30 days post-infection (PI), all animals were treated with one or two cycles of antibiotics as described in the materials and methods for 5 days, followed by a 3-5 day washout period. Mice with sterile urine were challenged with 10^7^ CFU of (**A, C**) UPEC strain UTI89-GFP-amp^R^ or (**B, C**) *E. faecalis* strain OG1RF_intergenicRS00490RS00495::Tn and sacrificed 24 hours post-challenge infection. Thirteen cytokines were measured in the supernatants of bladder homogenates by multiplex assay. Spider plots show the median values of absolute bladder cytokine levels (pg/mL) on a log-scale at 24 hours post-primary (dotted lines) or challenge infection (solid lines). DPI - days post-infection. Data are pooled from 2-7 experiments, with 2-6 mice per group in each experiment, values for cytokine expression in naïve and primary infected prostate are replotted from Figure 2 for ease in comparing primary to challenge infection. Significance was determined using the Mann-Whitney nonparametric test with correction for multiple testing to determine the false discovery rate (FDR) adjusted p value: q<0.05. All *p*- and *q*-values for each comparison are listed in **Tables 7-8**.

**Figure S3.**
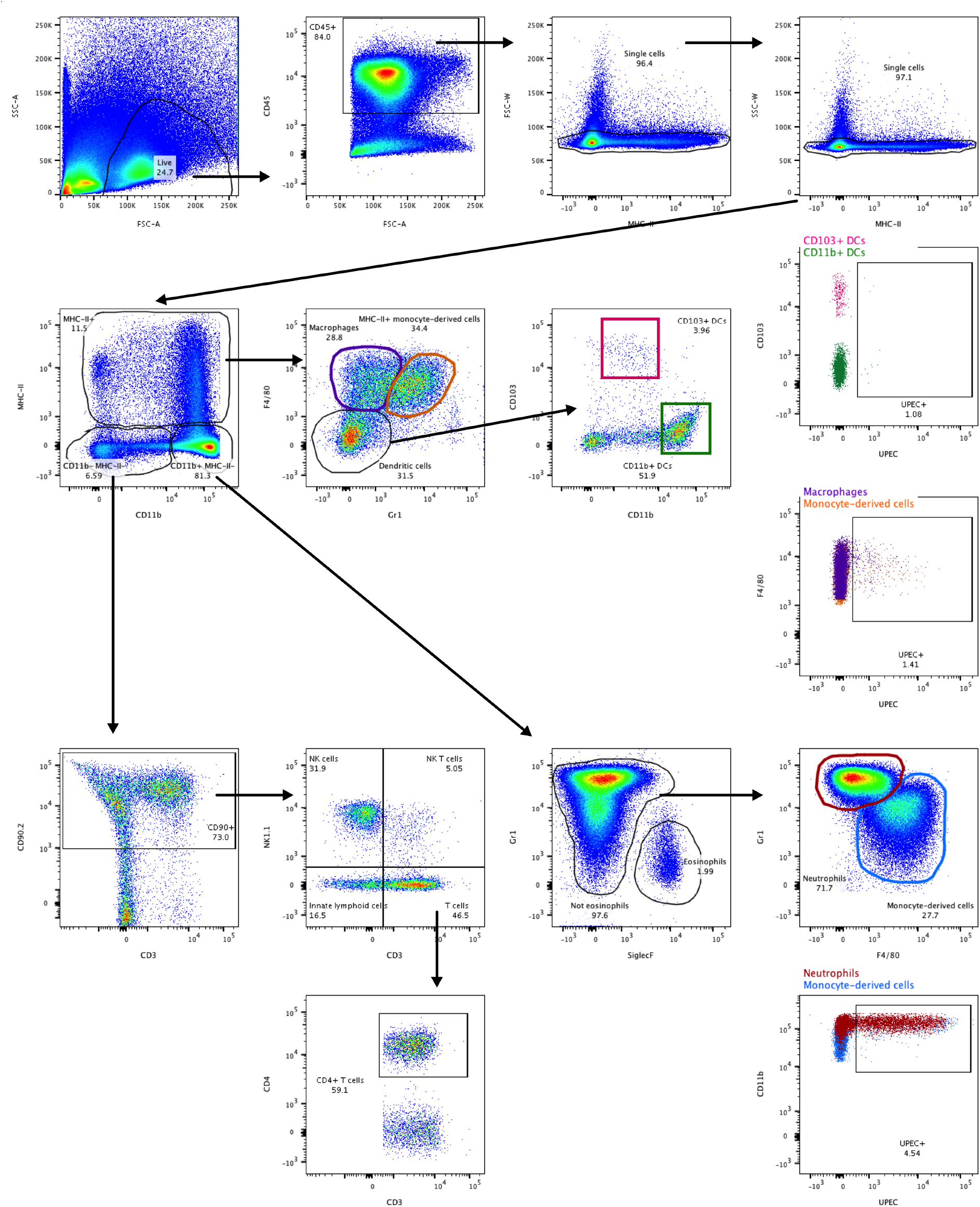
Gating strategy to identify prostate immune cells and bacteria^+^ immune cells. The plots show the gating strategies used for analyses illustrated in Figures 4, 5, and 6. 6-8-week old male C57BL/6 mice were instilled with 10^7^ CFU of either red fluorescent UPEC strain UTI89-RFP-kan^R^ or *E. faecalis* OG1RF stained with wheat germ agglutinin conjugated to Alexa Fluor 594. At 24 hours post-infection, mice were sacrificed, and prostates were digested and immunostained as described in the materials and methods and previously (Zychlinsky Scharff et al., 2019). Each sample, consisting of a single prostate, was acquired on an LSRFortessa SORP (BD Biosciences). Doublets were excluded from the CD45^+^ immune cell population. Macrophages were identified as MHC II^+^ CD11b^+^ F4/80^+^ CD64^+^ Ly6C/Ly6G^−^; dendritic cells (DCs) were MHCII^+^ CD11b^+/−^ CD103^+/−^ F4/80^−^ CD64^−^. Lymphocytes were gated as MHCII^−^ CD11b^−^ CD90.2^+^; then, NK cells were identified as NK1.1^+^ CD3^−^; NK T cells were NK1.1^+^ CD3^+^; T cells were NK1.1^−^ CD3^+^; CD4^+^ T cells were NK1.1^−^ CD3^+^ CD4^+^; innate lymphoid cells were NK1.1^−^ CD3^−^. Monocyte^−^derived cells were CD11b^+^ Ly6C/Ly6G^+/int^ MHCII^−^ or MHCII^+^ F4/80^int^ CD64^int^; neutrophils were CD11b^+^ Ly6C/Ly6G^+^ MHCII^−^ F4/80^−^ CD64^−^; eosinophils were CD11b^+^ SiglecF^+^ Ly6C/Ly6G^−/int^ MHCII^−^ F4/80^int^ CD64^−^. Immune cells engulfing bacteria were identified as specified above and then RFP^+^ cells were identified.

**Figure S4.**
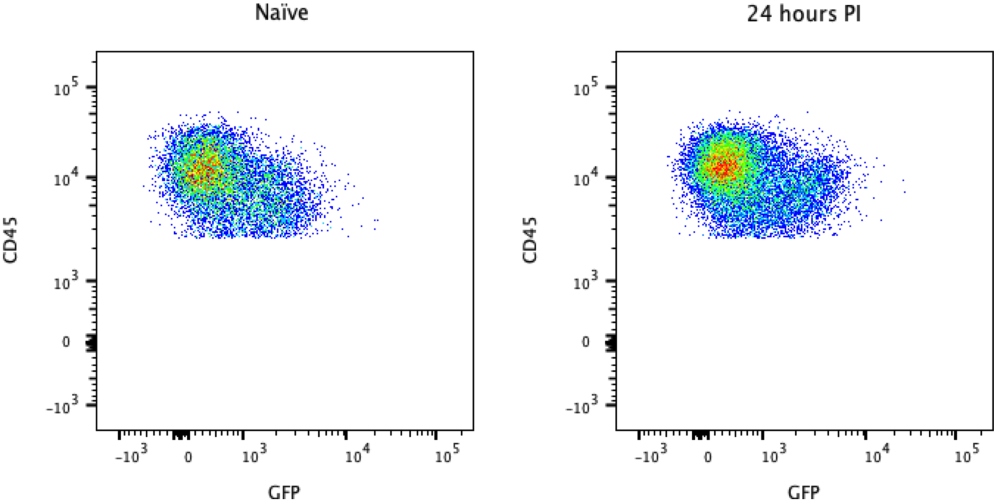
Prostate autofluorescence prevents detection of green-fluorescent OG1RF strain. 6-8-week old male C57BL/6 mice were instilled with 10^7^ CFU of either green fluorescent protein expressing OG1RF (OG1RF-GFP, 24 hours PI) or not infected (naïve). At 24 hours post-infection (PI), mice were sacrificed, and prostates were collected and processed for flow cytometry. Plots show CD45^+^ immune cells in the GFP channel from a naïve and an infected prostate following doublet exclusion.

